# Persistent cell motility requires transcriptional feedback of cytoskeletal — focal adhesion equilibrium by YAP/TAZ

**DOI:** 10.1101/265744

**Authors:** Devon E. Mason, James H. Dawahare, Trung Dung Nguyen, Yang Lin, Sherry L. Voytik-Harbin, Pinar Zorlutuna, Mervin C. Yoder, Joel D. Boerckel

**Author notes:** **Corresponding Author:** Joel D. Boerckel.

## Abstract

Cell migration initiates by traction generation through reciprocal actomyosin tension and focal adhesion reinforcement, but continued motility requires adaptive cytoskeletal remodeling and adhesion release. Here, we asked whether *de novo* gene expression contributes to this cytoskeletal feedback. We found that global inhibition of transcription or translation does not impair initial cell polarization or migration initiation, but causes eventual migratory arrest through excessive cytoskeletal tension and over-maturation of focal adhesions, tethering cells to their matrix. The transcriptional co-activators YAP and TAZ mediate this feedback response, modulating cell mechanics by limiting cytoskeletal and focal adhesion maturation to enable persistent cell motility and 3D vasculogenesis. Motile arrest after YAP/TAZ ablation was partially rescued by depletion of the YAP/TAZ-dependent myosin phosphatase regulator, NUAK2, or by inhibition of Rho-ROCK-myosin II. Together, these data establish a transcriptional feedback axis necessary to maintain a responsive cytoskeletal equilibrium and persistent migration.

## Introduction

Embryonic morphogenesis and post-natal regeneration rely on individual and collective cell migration to produce heterogeneous, organized, and vascularized tissues. Each cell is equipped with cytoskeletal and adhesion machinery that enable motility in response to physical cues communicated at both cell-cell and cell-matrix interfaces. Cell migration is driven by actin polymerization and actomyosin force generation, which coordinates focal adhesion formation, reinforcement, and disassembly^1–3^. These motile machinery form a molecular clutch, comprised of abundantly expressed proteins capable of producing intracellular tension, cellular polarization, and motility without *de novo* protein production, enabling rapid cellular responses to dynamic stimuli. However, the cytoskeletal activation that produces cell motility also induces mechanosensitive transcriptional programs. How new gene transcription regulates migration remains incompletely understood. Here, we sought to elucidate the role of transcriptional feedback in actomyosin control of cell migration.

Actomyosin tension is important for forward motility, but alone cannot not produce persistent migration, which requires coordinated actin treadmilling, leading-edge adhesion formation, and trailing-edge disassembly^4–6^. Thus, negative feedback systems are inherent to the migratory machinery. For instance, myosin phosphatases (e.g., MLCP) modulate motor activity to tune cytoskeletal tension^7–9^, while focal adhesion kinase (FAK) regulates adhesion remodeling to prevent cellular anchorage^10^.

The paralogous transcriptional co-activators yes-associated protein (YAP) and transcriptional co-activator with PDZ-binding motif (TAZ, also known as WWTR1) have recently emerged as important mechanotransducers that couple biophysical cell-cell and cell-matrix cues to mechanotransductive gene expression^11^. YAP/TAZ activity is regulated by subcellular localization, and their nuclear accumulation is induced by tension of the actomyosin cytoskeleton^11,12^. These observations position YAP and TAZ as potential key mediators of cytoskeleton-induced transcriptional programs.

ECFCs are blood-circulating endothelial cells^13^ that exhibit high proliferative and motile capacity and contribute to endothelium repair *in vivo*^14,15^. When cultured in three-dimensional matrices *in vitro* or transplanted *in vivo*, ECFCs have significant vasculogenic activity, characterized by cytoplasmic vacuolation, lumenization, and inosculation with host vasculature when implanted *in vivo*^16–18^. Here, we used ECFCs as a model system to demonstrate the importance of new gene transcription and translation for persistent cell migration and identify YAP and TAZ as mechanosensitive mediators of a transcriptional feedback loop that modulates cytoskeletal tension and focal adhesion formation. YAP and TAZ prevent myosin-dependent motile arrest by negatively regulating myosin light chain phosphorylation to enable persistent cell motility. Physiologically, we found that YAP and TAZ are essential for neovascular tube formation, 3D vacuolation, and neovascular sprouting.

## Results

### Transcription is essential for migration and regulates stress fiber and focal adhesion maturation

To evaluate directed cell motility driven by cell-cell and cell-matrix interactions in response to contact-inhibition release, we used the monolayer scratch assay on collagen-coated substrates to track migration over 8 hours (Figure 1a). To decouple existing cytoskeletal component function from *de novo* gene transcription/translation, we quantified longitudinal wound closure and wound migration rate in the presence of vehicle (DMSO) or inhibitors that prevent mRNA transcription (actinomycin D; 0.1 or 0.25 µg/mL) or protein translation (puromycin; 1 µg/mL), applied 1 hour prior to migration initiation (Figure 1a). In vehicle-treated cells, wound closure rate reached a plateau, or migratory equilibrium, in approximately 2 hours. Transcription inhibition significantly reduced wound closure percentage and rate by 8 hours after contact inhibition release (p < 0.0001, Figure 1b), while translation inhibition significantly slowed migration after 2 hours (p < 0.001), leading to motile arrest by 8 hours.

**Figure 1:**
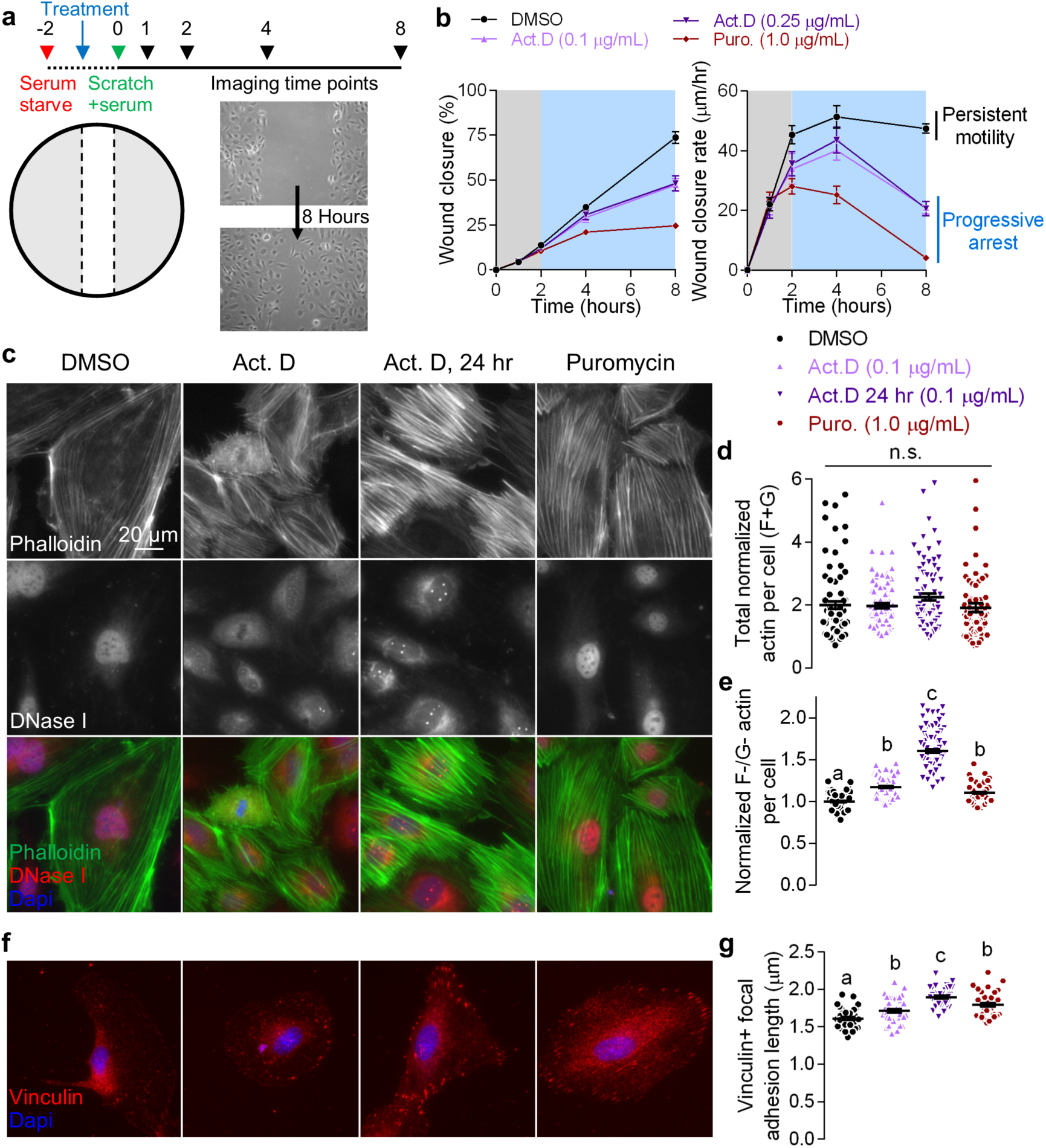
*De novo* protein synthesis is essential for actin cytoskeleton and focal adhesion dynamics during migration. ECFC migration after inhibition of *de novo* protein synthesis was assessed using the wound migration assay followed by immunofluorescence. **a**, Confluent ECFCs were serum starved for 2 hours and actinomycin D or puromycin were added 1 hour into the serum starve. Monolayers were scratched to form an open “wound” over which to longitudinally quantify migratory closure. **b**, Wound closure percentage, measured as (initial wound area – wound area at 8 hours)/initial wound area × 100, and wound closure rate, measured as the distance the cell front moved over each imaging period (µm/hr). Background color shows *de novo* gene expression-independent (grey) and –dependent (blue) phases. n = 19-24, p < 0.025, two-way ANOVA with Tukey’s *post hoc* test. **c,** Representative images of F- and G-actin visualized by Alex Fluor 488-conjugated phalloidin and Alexa Fluor 594-conjugated DNase I. Nuclei were visualized by DAPI. **d**, Total actin intensity (i.e., sum of normalized phalloidin and DNase I fluorescent intensity). **e**, F-/G-actin ratio per cell, normalized to DMSO-treated controls. n = 60 cells, p < 0.0001, Kruskal Wallace with Dunn’s *post hoc* test. **f**, Representative images of vinculin and nuclei visualized by Alexa Fluor 594-conjugated secondary and DAPI. **g,** Quantification of mature vinculin+ focal adhesion length. n = 40 cells, p < 0.009, ANOVA with Tukey’s *post hoc* test. Repeated significance indicator letters (e.g., a–a) signify p > 0.05, while groups with distinct indicators (a vs. b) signify p < 0.05. Summary statistics are represented as mean ± s.e.m.

One possible explanation for delayed loss of cell motility after transcription inhibition is that the cytoskeletal components of the molecular clutch become depleted over time. To test this, we evaluated actin polymerization and focal adhesion formation by immunofluorescence in leading-edge cells. Rather than observing depletion of actomyosin or focal adhesion proteins, transcription/translation inhibition significantly increased stress fiber formation (Figure 1c), actin polymerization (Figure 1d,e), and focal adhesion maturation (Figure 1f,g). Total actin was computed as the sum of F- and G-actin intensities, normalized to control cell intensities, on a per-cell basis (by phalloidin and DNaseI, respectively) and was unchanged by transcription or translation inhibition (p > 0.17; Figure 1d). Rather, the fraction of filamentous to globular actin (i.e., F/G-actin) was significantly increased by both inhibitors (Figure 1e). Vinculin recruitment to focal adhesions was increased by transcription or translation inhibition, significantly increasing focal adhesion length (Figure 1g). Prolonged treatment with actinomycin D (24 hrs) caused the most pronounced changes in actin and focal adhesion morphology.

Thus, rather than depleting the migratory machinery, inhibition of new gene expression led to abundant cytoskeletal tension and increased focal adhesions. This suggests a transcriptional feedback mechanism that is responsible for modulating cytoskeletal tension and focal adhesion dynamics to enable persistent migration.

### YAP and TAZ are mechanosensitive in ECFCs

Recent evidence has identified the transcriptional co-activators, YAP and TAZ, as cytoskeletal tension-activated regulators of gene expression^11^. Therefore, we hypothesized that YAP and TAZ mediate this transcriptional feedback. First, we confirmed that YAP and TAZ are mechanosensors of cytoskeletal tension in ECFCs using collagen-coated extracellular matrices of variable rigidity. Cells were seeded in sparse (8,500 cells/cm^2^) conditions on soft (1.85 kPa) or stiff (29 kPa) polyacrylamide (PA) or glass overnight followed by fixation and visualization of YAP and TAZ localization by immunocytochemistry (Supplementary Figure 1a). Consistent with prior reports in other cell types^11^, increased substrate rigidity significantly increased spread cell area and elongation (Supplementary Figure 1b,c) and increased nuclear localization of both YAP and TAZ (Supplementary Figure 1d).

### Dynamic control of YAP and TAZ promote migration and cytoskeletal pre-stress independent of angiocrine expression

YAP/TAZ subcellular localization is regulated by cell-cell interactions through the Hippo pathway^19^ and by cell-matrix interactions through cytoskeletal tension associated with cell spreading^11,12,20^.

To track these interactions during cell migration, we quantified cell density and area as a function of position relative to the original wound edge, 10 hours after migration initiation (Figure 2a,b). Cell density remained constant in the cell monolayer at distances greater than 500 µm behind the wound edge, but dropped exponentially (R^2^ = 0.90) to the leading edge (Figure 2a). Spread cell area similarly increased with an exponential fit (R^2^ = 0.68), but the effect of the wound on spread area extended only 200 µm into the monolayer. Total (Figure 2c) and nuclear (Figure 2d) YAP and TAZ fluorescent intensity increased preferentially in migrating cells beyond the original wound edge (Figure 2c, d). The increase in total YAP/TAZ levels could be explained either by increased expression or by stabilization from proteasomal degradation^19,21^. However, YAP/TAZ mRNA expression was not increased after contact inhibition release (Supplementary Figure 2a,b), suggesting that YAP and TAZ are stabilized from degradation in migrating cells.

**Figure 2:**
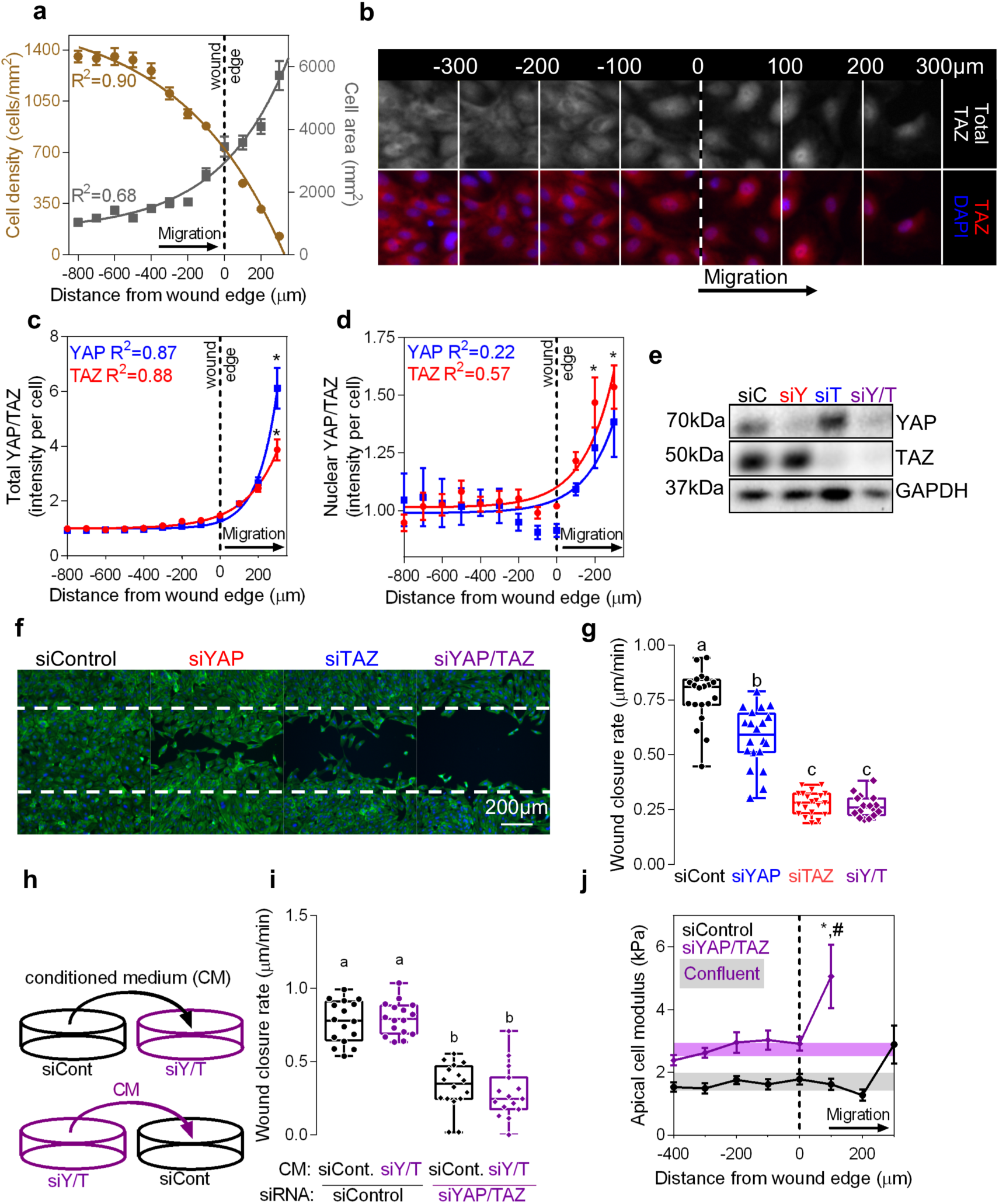
YAP and TAZ are essential for ECFC motility by limiting cytoskeletal pre-stress. Confluent ECFCs were scratched and imaged over 12 hours and fixed for immunofluorescence. **a,** Average cell density (*y* = −1863*e*^−.00177*x*^+ 1643) and area (*y* = 7062 − 6287(1 – *e*^−.00267*x*^)) as a function of distance from the leading edge (dotted lines) in 100 µm ROI’s. **b,** Representative immunofluorescent images of TAZ localization visualized by Alexa Fluor 594-conjugated secondary and DAPI subdivided into 100 µm ROI’s. **c,** Normalized total YAP (*y* = .255*e*^−.00996*x*^ + 1) and TAZ (*y* = .486*e*^−.00582*x*^ + 1) fluorescent intensity. **d,** Normalized nuclear YAP (*y* = .033*e*^−.00839*x*^ + 1) and TAZ (*y* = .120*e*^−.00529*x*^ + 1) fluorescent intensity. n = 7, * indicates p < 0.0001 vs all other ROI’s, two-way ANOVA with Tukey’s *post hoc* test. **e,** Representative (n = 3) immunoblot against YAP and TAZ in ECFCs treated with siRNA targeting YAP and/or TAZ. **f,** Representative images of ECFC migration after YAP and/or TAZ depletion, with actin visualized by Alexa Fluor 488-conjugated phalloidin. **g,** Wound closure rate. n = 16-20, p < 0.0001, ANOVA with Tukey’s *post hoc* test. **h,** Conditioned media swap experiment. Medium was conditioned overnight by siControl or siYAP/TAZ cells, then transferred to adjacent samples containing siControl or siYAP/TAZ cells for wound closure experiments. **i,** Wound closure after conditioned media treatment. n = 16, p < 0.0001, ANOVA with Tukey’s *post hoc* test. **j,** Apical cell modulus measured by nanoindentation at 100 µm regions of interest (ROI). n = 10-32 cells per ROI, * p < 0.023 vs control leading edge, # p < 0.001 vs siYAP/TAZ monolayer, two-way ANOVA with Tukey’s *post hoc* test. Repeated significance indicator letters signify p > 0.05, while groups with distinct indicators signify p < 0.05. Summary statistics in **a**, **c**, **d**, and **j** are represented as mean ± s.e.m.

To test the combinatorial roles of YAP/TAZ in cell motility, we depleted YAP and/or TAZ using RNAi, reducing protein expression to 27% and 18% of scrambled siRNA controls, respectively (Figure 2e; Supplementary Figure 2c,d). Control cells closed the 0.5mm-wide gaps within 12 hours, but YAP and/or TAZ depletion significantly impaired wound closure, with TAZ and YAP/TAZ depletion nearly abrogating migration (Figure 2f,g). YAP/TAZ depletion significantly reduced mRNA expression of secreted growth factors and enzymes including CTGF, Cyr61, and SerpinE1 (Supplementary Figure 2f,g). This suggested that YAP/TAZ activation could stimulate cell migration by induction of secreted angiocrines. However, recombinant reconstitution of these proteins failed to rescue cell motility (Supplementary Figure 2h,i) and transposition of conditioned medium from either control or YAP/TAZ-depleted cells (Figure 2h) similarly had no effect on either control or YAP/TAZ-depleted cell motility (p = 0.99; Figure 2i), suggesting a cell-intrinsic mechanism. Additionally, inhibition of proliferation using the DNA cross-linking agent, mitomycin C (mito. C), had no effect on wound closure in either control or YAP/TAZ-depleted cells (p > .30; Supplementary Figure 2j).

Intracellular mechanics dynamically respond to extracellular stimuli like contact inhibition release during migration^22–24^ and drive YAP/TAZ nuclear localization^11,20^. Cellular resistance to perpendicular compressive forces at the cell apex is mediated by the tensile actomyosin network. The collective stress imposed by myosin motors on bundled actin fibers is referred to as cellular pre-stress, and can be quantified as apical cell modulus, measured by single-cell nanoindentation.^22^ We measured apical cell modulus as a function of migratory distance (Figure 2j). In control cells, apical modulus was elevated at the leading edge of the migrating front (Figure 2j), consistent with tension-induced YAP/TAZ activation (cf. Figure 2d). YAP/TAZ-depleted cells exhibited reduced migration as above, but featured significantly elevated cell modulus compared to controls, both in the monolayer (p = 0.005) and at the leading edge (p = 0.001; Figure 2j).

### YAP and TAZ enable ECFC migration and directional persistence

Next, by live cell imaging, we evaluated YAP/TAZ function in the motility of single cells during collective and individual cell migration (Figure 3; Supplementary Videos 1-3). Control cells exhibited persistent, directional collective migration, whereas YAP/TAZ-depleted cells remained tethered in place, with many cells exhibiting small displacement between imaging intervals (15 min) but without persistent motion beyond the original cell borders (Figure 3a). Instantaneous, scalar, cell migration speed was evaluated on a single-cell basis both for cells at the leading edge and in trailing cells, whose motility depends on contact inhibition release by motion of the leading cells. Control cell migration speed (in both leading and trailing cells) peaked 15 minutes after wounding, and decreased to a minimum after 2 hours (Figure 3b) before re-accelerating until experiment completion at 10 hrs. In contrast, YAP/TAZ-depleted cell migration speed was initially lower and then decreased continuously until hour 10. Average migration speed was equivalent in leading and trailing cells in both control and YAP/TAZ-depleted conditions; however, YAP/TAZ-depletion significantly slowed migration regardless of position (Figure 3c).

**Figure 3:**
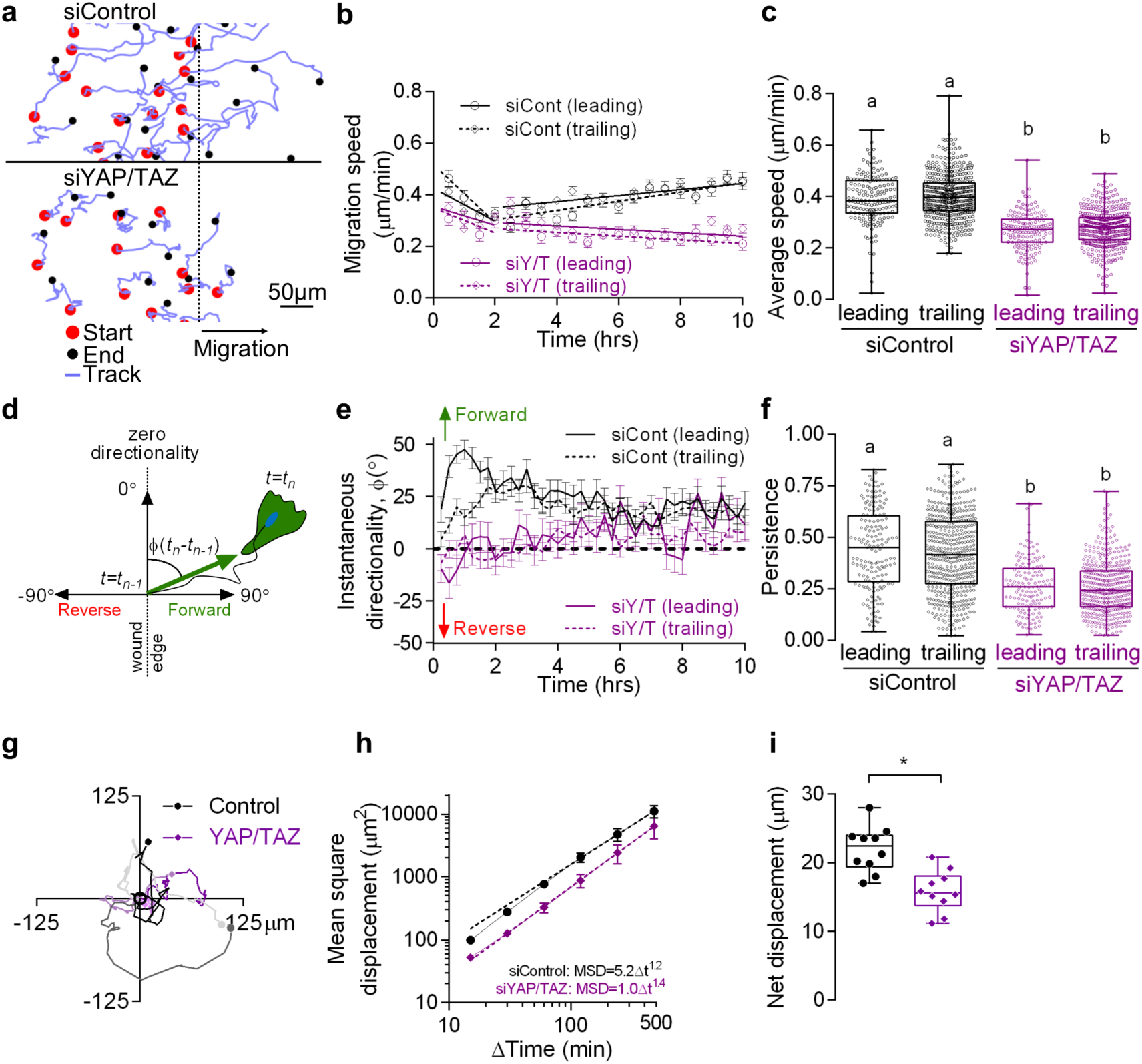
YAP and TAZ promote collective and individual cell motility. ECFCs expressing EGFP (siControl) or mTomato (siYAP/TAZ) were plated in wells of a custom PDMS stencil and imaged after the stencil was removed at 15-minute intervals over 10 hours to track individual cell migration. **a,** Representative cell migration tracks over 10 hours (dashed line indicates starting position). **b,** Instantaneous average cell migration speed (30-minute intervals shown) as a function of time. Cells were grouped into leading (< 100 µm from the front most cell) and trailing (100-500 µm from the front most cell) based on their initial position. n = 140-165 leading cells and n = 437-484 trailing cells. **c,** Average migration speed per cell, averaged over 10 hours. p < 0.0001, ANOVA with Tukey’s *post hoc* test. **d**, Schematic of instantaneous cell directionality, defined as the direction, *⇕*(*t*_*n*_-*t*_*n*-1_), a cell moved between the current position, *t*_*n*_, and the previous position, *t*_*n*-1_, relative to the wound edge. **e**, Instantaneous cell directionality in leading and trailing cells. Data points above 0° indicate cells in that group tend to move into the wound edge whereas those below tend to move in the direction opposite the wound edge. **f,** Motile persistence, quantified as end-to-end displacement divided by total displacement. p < 0.0001, ANOVA with Tukey’s *post hoc* test. **g,** Random migration of mTomato-expressing siYAP/TAZ cells mixed with a 100-fold excess of EGFP-expressing siControl cells. **h,** Mean square displacement of randomly migrating siControl (R^2^ = 0.57) and siYAP/TAZ (R^2^ = 0.35) cells. **i,** Average net displacement of randomly migrating cells over 10 hours. n = 10, p < 0.0004, two-tailed Student’s unpaired t-test. Repeated significance indicator letters signify p > 0.05, while groups with distinct indicators signify p < 0.05. Summary statistics in **b, e**, and **h** are mean ± s.e.m.

Wound repair and directed angiogenesis *in vivo* require not only cell movement, but directional migration^25^. We therefore quantified the instantaneous directionality of individual cells both at the leading edge and in the trailing monolayer. Directionality was defined for each 15 minute time interval as the angle, *⇕*(*t*_*n*_-*t*_*n*-1_), relative to the wound edge, between an individual cell position at time *t*_*n*_ and its prior position at time *t*_*n*-1_ (Figure 3d). Leading edge control cells initiated forward migration, while trailing cells lagged by 2 hours, coinciding with the inflection in migration speed (cf. Figure 3b) and the time at which transcription inhibition begins to reduce cell motility (cf. Figure 1b). In contrast, YAP/TAZ-depleted cells exhibited zero average directionality (Figure 3e) and had reduced persistence (p < 0.0001), defined as the ratio of net migration distance to total distance traveled (Figure 3f).

We next asked whether the motility of individual YAP/TAZ-depleted cells could be restored by contact with control cells. YAP/TAZ-depleted cells were plated sparsely in a more than 100-fold excess of control cells and random migration within the mixed monolayer was tracked over time in 2D space (Figure 3g). YAP/TAZ-depletion significantly reduced individual cell motility by 43-47%, measured by mean square displacement (Figure 3h) and decreased net displacement (Figure 3i) compared to control cells in the same mixed monolayer. These data further support a cell-autonomous role of YAP and TAZ in cell migration.

### YAP and TAZ are dispensable for microtubule polarization and Golgi reorientation

Cell migration initiates by establishment of front-rear cell polarity, determining motile direction^4^. This cellular polarization requires microtubule-mediated polarization of the microtubule organizing center (MTOC) to coordinate microtubule extension and cytoskeletal remodeling, and is accompanied by polarization of the Golgi apparatus^26^. To determine whether YAP and TAZ regulate migratory cell polarity, we evaluated microtubule network structure and Golgi polarization, defined as orientation of the Golgi apparatus within ±60° of the direction of the wound (Figure 4a). We found that both control and YAP/TAZ-depleted cells had similar microtubule networks that extended from the MTOC to the cell periphery (Supplementary Figure 3a), and YAP/TAZ depletion had no effect on Golgi polarization in leading edge cells (Figure 4b). In cells at the migratory front, 60% of cell Golgi polarized in each condition, significantly greater than the random (33%) distribution observed in non-wounded monolayers (p < 0.0001; Supplementary Figure 3b). In the trailing region (Figure 4c), control cells also exhibited significant Golgi polarization (p = 0.002 vs random), but YAP/TAZ-depleted trailing cells did not (p = 0.81; Supplementary Figure 3b). These data suggest that YAP/TAZ are dispensable for direction sensing and the initiation of motile cell polarization, but are required for directional persistence.

**Figure 4:**
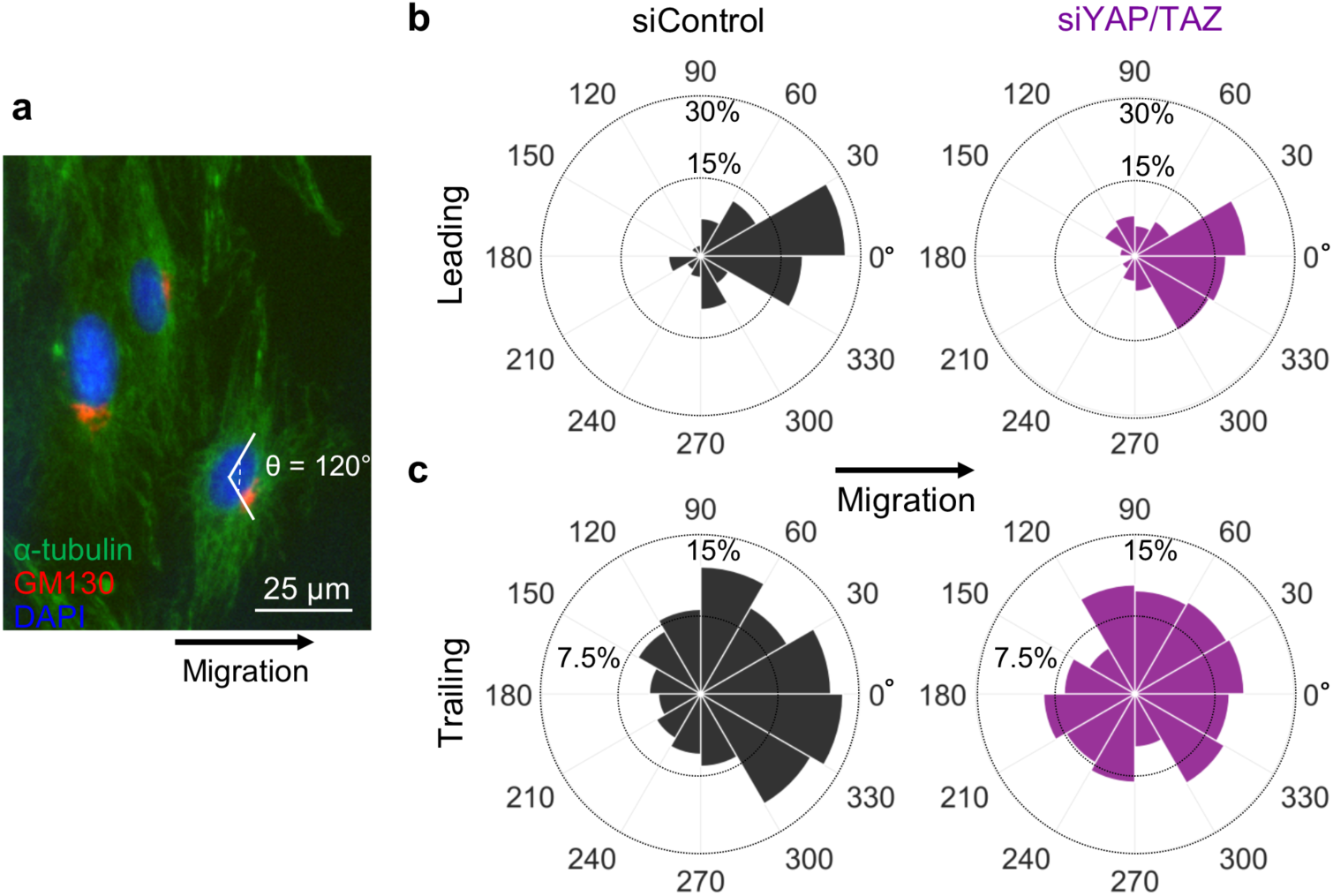
YAP and TAZ are dispensable for microtubule polarization. ECFCs were fixed 8 hours after the initiation of migration and immunofluorescence used to visualize microtubule polarization. **a,** Representative image of microtubules (α-tubulin) and Golgi (GM130) visualized with Alexa Fluor 488- and 594-conjugated secondary, respectively. Golgi were considered polarized when within a 120° region centered about a vector extending horizontally from the nuclei (DAPI) in the direction of the wound edge. **b,c,** Rose plot of Golgi polarization in siControl (black) and siYAP/TAZ (purple) in leading (**b)** and trailing (**c)** cells. n = 64-65 leading cells, n = 143-165 trailing cells.

### YAP and TAZ regulate cytoskeletal and focal adhesion remodeling

We found above that transcription inhibition caused migratory arrest and increased cytoskeletal polymerization and focal adhesion formation. Similarly, YAP/TAZ depletion impaired persistent migration with increased cellular pre-stress. We therefore hypothesized that YAP and TAZ transcriptionally regulate actin cytoskeletal architecture to enable persistent motility.

Consistent with this hypothesis, YAP and/or TAZ depletion from migrating cells significantly increased F-actin intensity and produced larger stress fibers (Figure 5a,b). While the total actin per cell (F+G) remained constant across conditions (p > 0.41; Figure 5d), YAP and/or TAZ depletion significantly increased F-actin polymerization from G-actin (Figure 5e).

**Figure 5:**
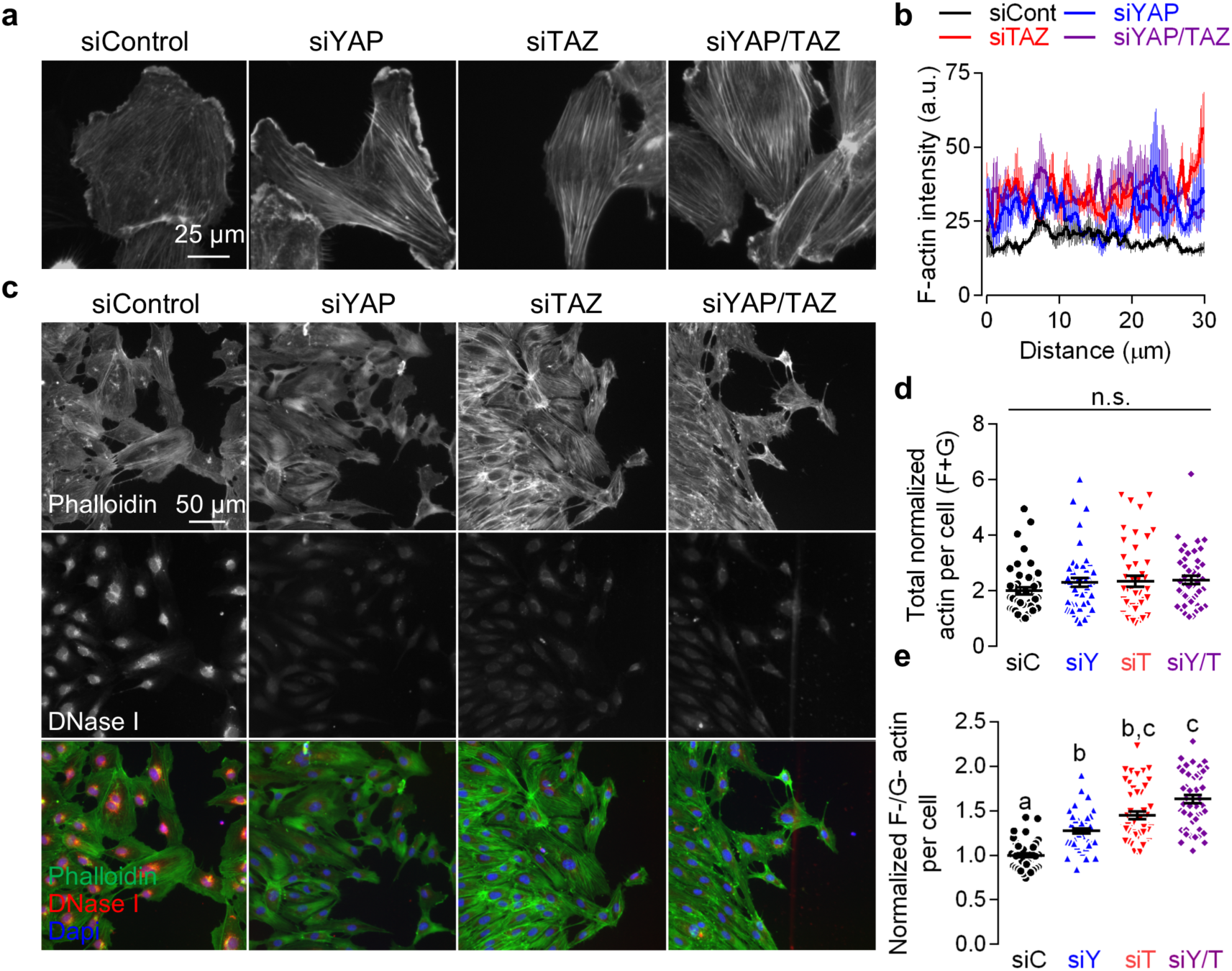
YAP and TAZ modulate actin polymerization and stress fiber formation. Migrating ECFCs were fixed 8 hours after the initiation of migration and actin was visualized using immunofluorescence. **a,** F-actin, visualized with Alexa Fluor 488-conjugated phalloidin. **b**, Average fluorescent intensity line profiles of phalloidin, line plots are the average of three cells per condition. **c,** Representative images of F- and G-actin visualized by Alex Fluor 488-conjugated phalloidin and Alexa Fluor 594-conjugated DNase I, respectively. **d,** Total actin intensity (i.e., sum of normalized phalloidin and DNase I fluorescent intensity). **e,** F-/G-actin ratio per cell, normalized to siControl. n = 45-50, p < 0.0001, ANOVA with Tukey’s *post hoc* test. Repeated significance indicator letters signify p > 0.05, while groups with distinct indicators signify p < 0.05. Summary statistics represented as mean ± s.e.m.

Cell motility requires formation of new focal adhesions at the cell’s leading edge and coordinated adhesion disassembly at the trailing edge. To visualize focal adhesions, we immunostained migrating cells for the focal adhesion protein, vinculin (Figure 6a). YAP/TAZ depletion increased the total number of vinculin+ focal adhesions (p < 0.0001; Figure 6b, left), and simultaneously reduced cell area and increasing cell elongation (Figure 6b, right, Supplementary Figure 4a-c).

**Figure 6:**
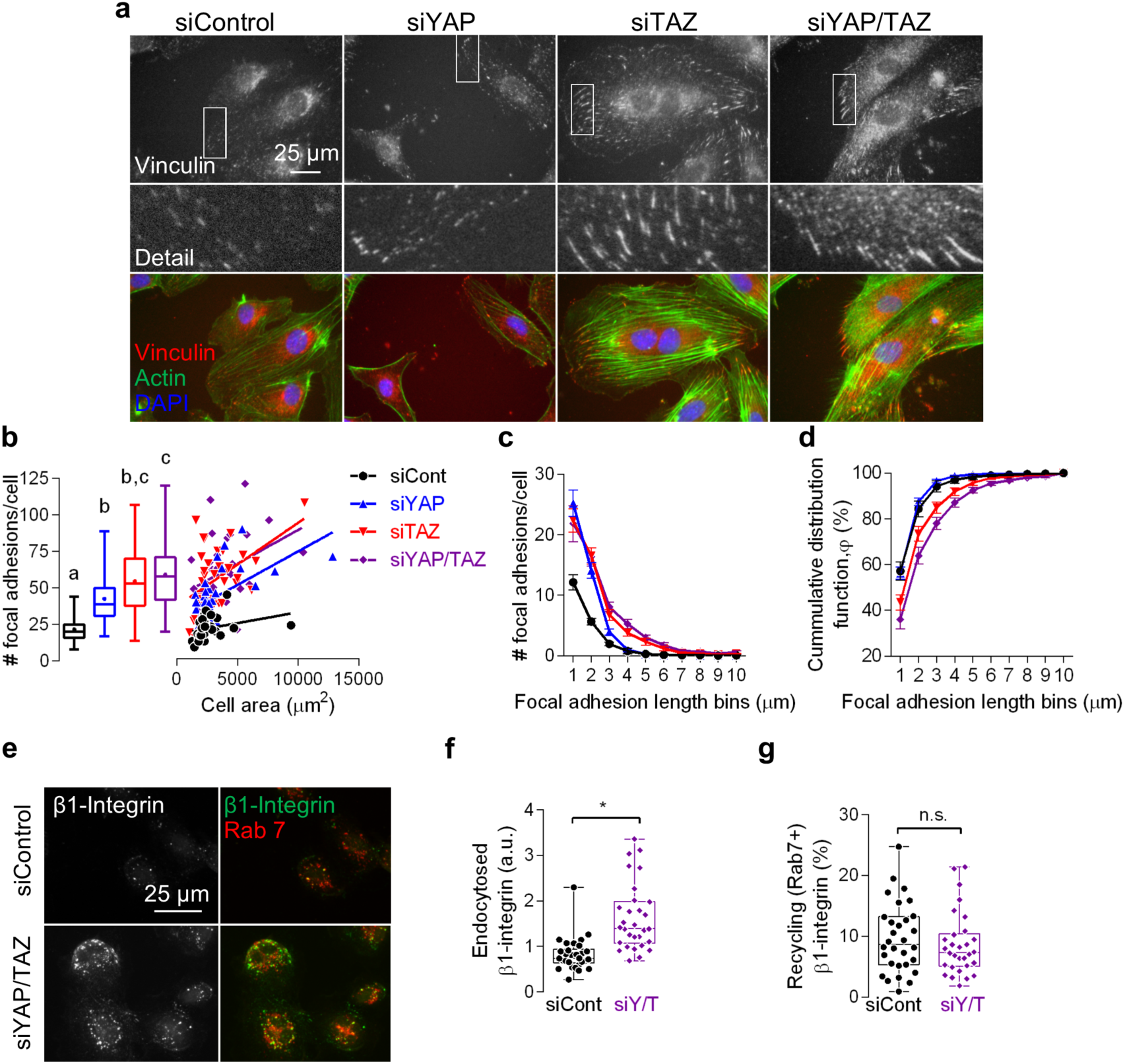
YAP and TAZ modulate focal adhesion remodeling without affecting β1-integrin recycling. Confluent ECFCs were fixed for immunofluorescence 12 hours after the initiation of migration. **a,** Representative images of vinculin and F-actin, visualized with Alexa Fluor 594-conjugated secondary and 488-conjugated phalloidin, respectively. **b,** Average number of focal adhesions per cell as a function of cell area. n = 30-34, p < 0.004, ANOVA with Tukey’s *post hoc* test. **c,** Focal adhesion size histogram and **d,** cumulative distribution function, representing the percentage of focal adhesions in a cell of a given size. **e,** Visualization of β1-integrin internalization and recyling. Live ECFCs were incubated with mouse monoclonal antibodies targeting active β1-integrin (10 µg/mL), which was endocytosed for 45 minutes, followed by acid wash and fixation. Internalized integrin was detected with Alexa Fluor 488-conjugated secondary. Rab 7+ endosomes were visualized with Alexa Fluor 594-conjugated secondary. **f,** Total fluorescent intensity of endocytosed β1-integrin. n = 30, p < 0.0001, two-tailed Student’s unpaired t-test. **g,** Fluorescent intensity of recycling endocytosed β1-integrin co-localized with Rab 7+ endosomes. p > 0.46, two-tailed Student’s unpaired t-test. Repeated significance indicator letters signify p > 0.05, while groups with distinct indicators signify p < 0.05. Summary statistics in **c** and **d** are mean ± s.e.m.

Focal adhesions enlarge and lengthen as they mature^27^. To test the effects of YAP/TAZ depletion on focal adhesion maturity, we quantified the number of focal adhesions in 1 to 10 µm length bins (Figure 6c,d). YAP depletion proportionally increased focal adhesion number regardless of length, while TAZ and YAP/TAZ depletion shifted the distribution to larger focal adhesions (Figure 6d), resulting in increased average focal adhesion length (Supplementary Figure 4d). Focal adhesion tyrosine kinase (FAK), was present and phosphorylated at tyrosine 397 (pFAK) in focal adhesions found in both control and YAP/TAZ depleted cells, but was preferentially localized to the cell periphery in YAP/TAZ depleted cells (Supplementary Figure 5).

### YAP and TAZ regulate focal adhesion formation and maturation, but do not inhibit adhesion disassembly

Observation of live cell migration revealed persistent actin-focal adhesion connectivity at the cell trailing edge, resulting in protrusion of actin stress fibers beyond the trailing cell membrane, tethering the cell at the training edge (Supplementary Videos 4, 5), similar to the integrin-bound tubular membrane microaggregates described in keratinocyte migration^28^. This led us to ask whether YAP/TAZ could regulate focal adhesion release or disassembly. To test this, we quantified internalization of the pro-angiogenic^29^ integrin, β1, using an active β1-integrin antibody internalization assay and evaluated β1 recycling by colocalization with Rab 7+ endosomes^30^ (Figure 6e). If YAP/TAZ control focal adhesion disassembly, we would expect to observe impaired integrin endocytosis with YAP/TAZ depletion. However, we found that β1-integrin endocytosis was significantly increased in YAP/TAZ depleted cells (p < 0.0001, Figure 6f), concomitant with increased focal adhesion number (cf. Figure 6a-d). Further, upon internalization, the amount of β1-integrin recycling in rab7+ endosomes was constant (p = 0.69, Figure 6g), suggesting that YAP/TAZ modulate focal adhesion formation, but not disassembly or recycling.

### YAP and TAZ limit cytoskeletal tension through MYL phosphorylation

Cytoskeletal tension and aggregation of actin filaments into stress fibers is mediated by myosin motor force generation and crosslinking, which in turn stabilizes focal adhesions^31^. Therefore, we next asked whether YAP and TAZ regulate cytoskeletal remodeling through activation of non-muscle myosin II. We found that YAP/TAZ depletion increased serine 19 phosphorylation of myosin light chain (pMYL), which localized to stress fibers, consistent with mechanosensitive recruitment of myosin II to regions of high actin stress^32,33^ (Figure 7a). YAP/TAZ depletion did not decrease the total amount of MYL (p = 0.15), but significantly increased the amount and percentage of pMYL (p < 0.0001, Figure 7b,c).

**Figure 7:**
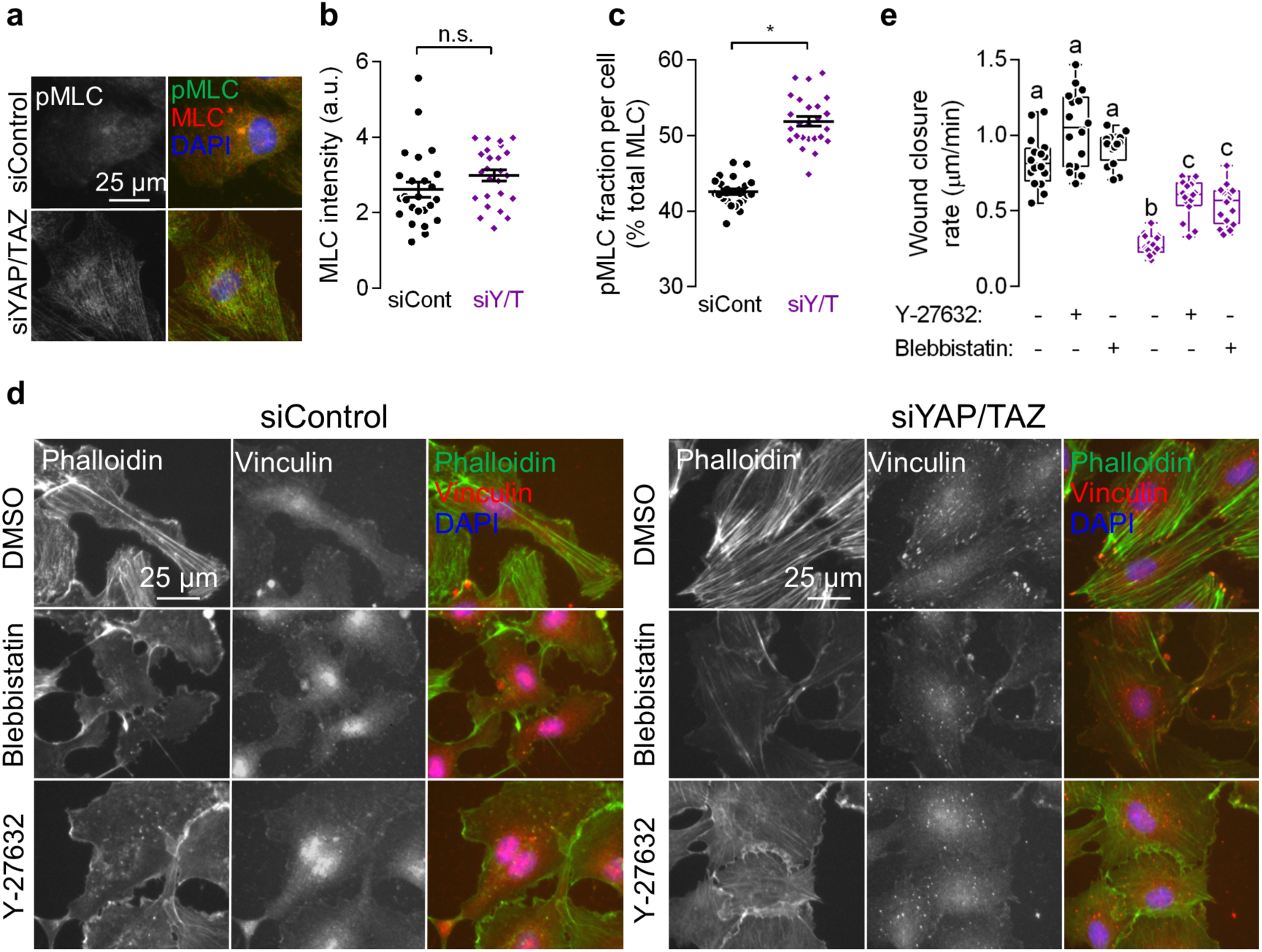
YAP and TAZ promote migration by limiting myosin light chain phosphorylation. Migrating ECFCs were treated with ROCK (Y-27632) or non-muscle myosin II (Blebbistatin) inhibitors and imaged over 12 hours then fixed for immunofluorescence. **a,** Representative images of MLC and pMLC visualized by Alexa Fluor 594- and 488-conjugated secondary, respectively. **b,** Total MLC intensity per cell. n = 25, p > 0.15, two-tailed unpaired Student’s t-test. **c,** Percentage of MLC phosphorylated at Ser19 (i.e., pMLC intensity / total MLC intensity × 100, per cell). n = 16, p < .0001, two-tailed unpaired Student’s t-test. **d,** Vinculin and actin in migrating ECFCs visualized by 594-conjugated secondary and 488-conjugated phalloidin. **e,** Wound closure rates after treatment with Y-27632 (10 µM) or Blebbistatin (20 µM). n = 16, p < 0.0001, ANOVA with Tukey’s *post hoc* test. Repeated significance indicator letters signify p > 0.05, while groups with distinct indicators signify p < 0.05. Summary statistics in **b** and **c** are represented as mean ± s.e.m.

To confirm a functional role for myosin in YAP/TAZ-dependent cytoskeletal dynamics and migration, we inhibited myosin II ADP cycling using Blebbistatin and myosin association with actin by inhibiting Rho-associated kinase (ROCK)-mediated phosphorylation of MYL, using Y-27632. Both myosin and ROCK inhibition in YAP/TAZ-depleted cells reduced stress fiber formation and focal adhesion maturation to levels observed in control cells (Figure 7d) and substantially rescued cell migration (Figure 7e).

### YAP/TAZ regulate NUAK2 to control cytoskeletal polymerization

These data suggest that YAP/TAZ mediate feedback control of cytoskeletal and focal adhesion dynamics through the Rho-ROCK-myosin II pathway. We performed a meta-analysis of previously published ChIP-seq and gene expression data and identified SNF-like kinase 2 (NUAK2), Rho GTPase activating protein 28 (ARHGAP28), and ARHGAP29 as YAP/TAZ-dependent target genes^34–37^. We confirmed that YAP/TAZ regulate these putative targets by PCR of ECFC mRNA 1 hour after initiation of the wound assay. Both NUAK2 and ARHGAP28 were significantly increased after YAP/TAZ depletion (p < .0001), but ARHGAP29 was significantly reduced (p < .0001; Supplementary Figure 6a). These data confirm that YAP/TAZ regulate expression of cytoskeletal regulators. Notably, NUAK2 was upregulated 7-fold in YAP/TAZ-depleted cells. NUAK2 expression is induced by cytoskeletal tension to phosphorylate and deactivate myosin light chain phosphatase (MLCP) and sequester the myosin binding subunit (MYPT1) of MLCP preventing dephosphorylation of myosin II, negating ROCK-mediated myosin activation^8,9^. We therefore chose to orthogonally probe the function of myosin II phosphorylation by perturbing NUAK2 expression. These data implicate migration-inducible NUAK2 expression and function in YAP/TAZ-Rho-ROCK-myosin II-mediated cytoskeletal and focal adhesion feedback.

To better understand the kinetics of YAP/TAZ-dependent gene induction, we examined gene expression at 0, 1, and 12 hours after initiation of migration. As expected, YAP/TAZ depletion abrogated inducible expression of the canonical YAP/TAZ-TEAD target genes, Cyr61 and CTGF (Figure 8a; Supplementary Figure 6b), but also significantly up-regulated NUAK2 at 1 hour (Figure 8b), consistent with increased myosin phosphorylation (cf. Figure 7). We next tested whether YAP/TAZ/NUAK2 co-depletion (Supplementary Figure 6c) could restore actomyosin architecture compared to control cells. NUAK2 depletion alone did not alter total actin (p > 0.11, Figure 8c,d) and had no effect on F-/G-actin ratio in YAP/TAZ-intact cells, but significantly rescued the increase in actin polymerization caused by YAP/TAZ depletion (Figure 8e). We orthogonally validated these results by treating YAP/TAZ-depleted cells with WZ4003, a selective inhibitor of NUAK1/2^38^ (Supplementary Figure s7a). WZ4003 treatment significantly decreased actin polymerization as well as total actin in YAP/TAZ depleted cells (Supplementary Figure s7b,c).

**Figure 8:**
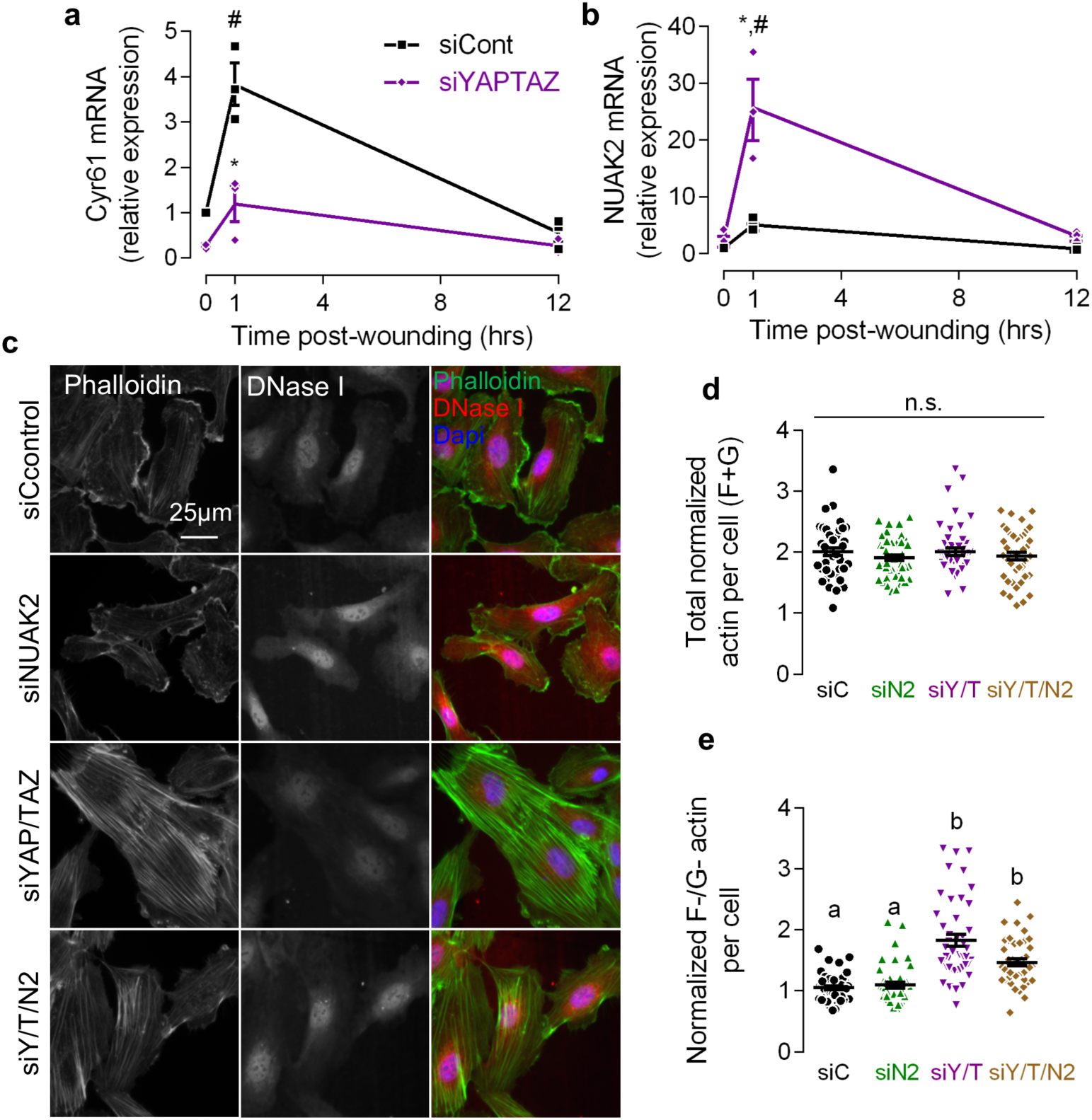
YAP and TAZ modulate actin polymerization through NUAK2 regulation. Migrating ECFC lysate was collected at 0, 1, and 12 hours after initiation of migration and gene expression analyzed by RT-qPCR. **a**, Cyr 61 and **b**, NUAK2 gene expression in migrating ECFCs. n = 3, * p < 0.0004 vs. control at 1 hour, # p < 0.0001 vs. 0 hours, two-way ANOVA with Tukey’s *post hoc* test. **c,** Representative images of F- and G-actin visualized with Alexa Fluor 488-conjugated secondary and 594-conjugated phalloidin. **d,** Total actin intensity measured as the sum of F- and G-actin intensity. n = 45 cells, p > 0.11, ANOVA with Tukey’s *post hoc* test. **e**, F-/G-actin ratio per cell, normalized to DMSO-treated controls. p < 0.005, ANOVA with Tukey’s *post hoc* test. Repeated significance indicator letters signify p > 0.05, while groups with distinct indicators signify p < 0.05. Summary statistics are represented as mean ± s.e.m.

### YAP and TAZ spatially control vinculin incorporation into structural focal adhesions via NUAK2

Together, these data implicate YAP and TAZ in feedback control of actomyosin tension through Rho-ROCK-myosin II to prevent cellular tethering at focal adhesions. To specifically evaluate this feedback in structural focal adhesions, we next used detergent solubilization during fixation to remove poorly adherent focal adhesions and Triton-soluble cytoskeletal elements, leaving the structural fraction^39^. The composition of the structural focal adhesions was determined by immunostaining for tension-dependent vinculin and tension-independent paxillin^27^ incorporation into focal adhesions in single cells (Figure 9a). YAP/TAZ depletion increased the amount of structural vinculin, without affecting structural paxillin (Supplementary Figure 8). Focal adhesion size, a measure of maturity, was increased by YAP/TAZ depletion, but NUAK2 co-depletion significantly rescued vinculin+ focal adhesion size (Figure 9c).

**Figure 9:**
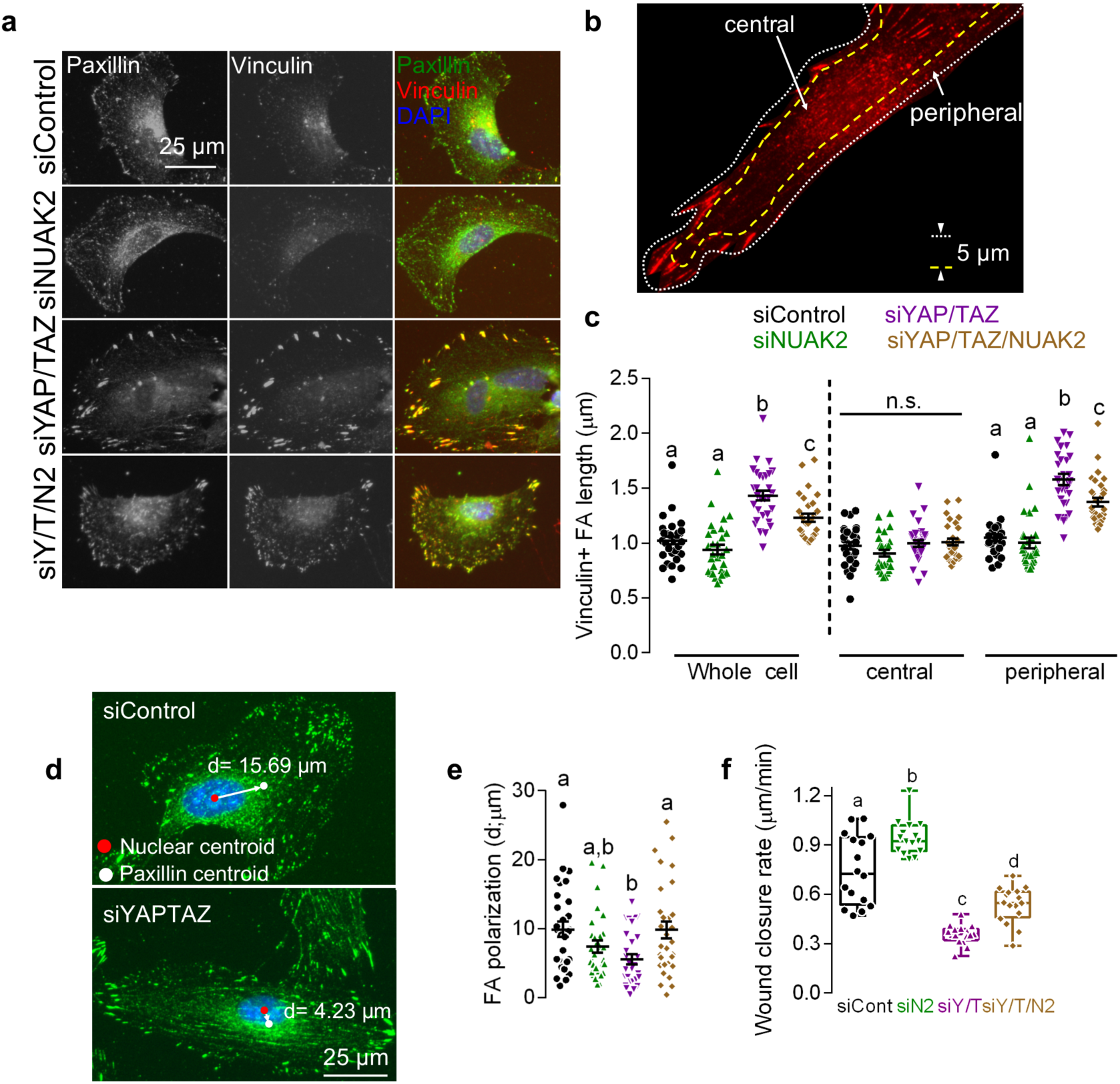
YAP/TAZ-regulated NUAK2 enhances focal adhesion maturation and polarization, resulting in reduced cell motility. ECFCs depleted of YAP/TAZ and/or NUAK2 were plated on collagen coated glass coverslips for 24 hours then triton-extracted concurrent with fixation for immunofluorescence and analysis of triton-insoluble structural focal adhesions. **a,** Representative images of vinculin and paxillin visualized with 594- and 488-conjugated secondary, respectively. **b,** Individual cell compartments were subdivided into peripheral and central (5 µm from the edge of the cell) regions to detect spatial differences in focal adhesion morphology. Focal adhesion length, an indicator of maturation, and number, an indicator of formation, where found in individual cells at both peripheral and central regions. **c,** Vinculin+ focal adhesion length in the whole cell (p < .003), central region (p > 0.1), and peripheral region (p < 0.006), ANOVA with Tukey’s *post hoc* test for the for whole cell and central region and Kruskal-Wallis with Dunn’s *post hoc* test for the peripheral region. **d,** Representative image of focal adhesion polarization distance indicated by the white line between the nucleus (red dot) and focal adhesion (white dot) intensity centroid. **e,** Focal adhesion polarization distance. p < 0.02, ANOVA with Tukey’s *post hoc* test. **f,** Confluent ECFCs were scratched and imaged over 12 hours and wound closure rate quantified, n = 16, p < 0.006, ANOVA with Tukey’s *post hoc* test. Summary statistics in **c and e** are represented as mean ± s.e.m. Repeated significance indicator letters signify p > 0.05, while groups with distinct indicators signify p < 0.05.

Focal adhesion remodeling and cell-generated traction is spatially controlled and occurs predominantly at leading and trailing edges^7,40^. Further, in YAP/TAZ-depleted cells, we observe preferential focal adhesion maturation at the cell periphery. Therefore, we subdivided focal adhesions in each cell into peripheral (5 µm from every edge) or central regions (Figure 9b). YAP/TAZ and YAP/TAZ/NUAK2 depletion had no effect on focal adhesion number or morphology within the central region of interest (Figure 9c, Supplementary Figure 8b); however, in the peripheral region, NUAK2 co-depletion significantly rescued focal adhesion length (Figure 9c). We further confirmed these observations using the NUAK1/2 inhibitor WZ4003, which normalized focal adhesion morphology in YAP/TAZ-depleted cells (Supplementary Figure 7d). Interestingly, WZ4003 caused near complete loss of vinculin incorporation into focal adhesions, more so than NAUK2 depletion or WZ4003 treatment in control cells.

### YAP/TAZ modulate myosin tension to enable focal adhesion polarization

Above, we observed that YAP/TAZ depletion impaired persistent and directional cell migration, but did not affect MTOC polarization or initial direction sensation. Persistent forward motility requires that focal adhesions preferentially form at the leading edge, mature in the lamellum, and disengage at the trailing edge^6,40^. To test whether YAP/TAZ regulate focal adhesion polarization, we quantified the distance between the centroid of the nucleus and the centroid of structural focal adhesions (Figure 9d). YAP/TAZ depletion significantly reduced focal adhesion polarization, which were fully restored by co-depletion of NUAK2 (Figure 9e). Finally, we found that NUAK2 co-depletion partially rescued YAP/TAZ-dependent anchorage release and migration (Figure 9f), but WZ4003 treatment did not (Supplementary Figure 7e).

### YAP and TAZ are essential for endothelial vacuole formation, tubulogenesis, and sprouting angiogenesis

ECFC vacuole formation is senstive to 3D matrix mechanical properties^16^ and early embryonic vascular remodeling requires hemodynamic force *in vivo*^41^. To control mechanical cues in 3D, we cultured ECFCs in collagen matrices of variable physiologic elasticity. Matrix rigidity was tuned, independent of collagen density, by modulating monomer:oligomer collagen ratio^16,17^ (Figure 10a,b). ECFCs were cultured in monomeric (G’ = 310 Pa / E = 9 kPa) or oligomeric collagen matrices (G’ = 1050 Pa / E = 23 kPa) whose rigidity approximated physiologic and pathologic extracellular matrix, respectively^17^ (Figure 10a). Stiffer oligomeric matrices qualitatively enhanced ECFC spreading, vacuolization, and network formation (Figure 10b), consistent with prior findings^16,17^. YAP and TAZ depletion completely abrogated 3D vacuolation and interconnected vasculogenic network formation in oligomeric collagen matrices (G’ = 132 Pa) (Figure 10c). Similarly, in the matrigel tubulogenesis assay, YAP and/or TAZ depletion combinatorially reduced tubular network length and number (Figure 10d,e), with a greater effect of TAZ vs. YAP depletion, consistent with their effects on cytoskeletal dynamics.

**Figure 10:**
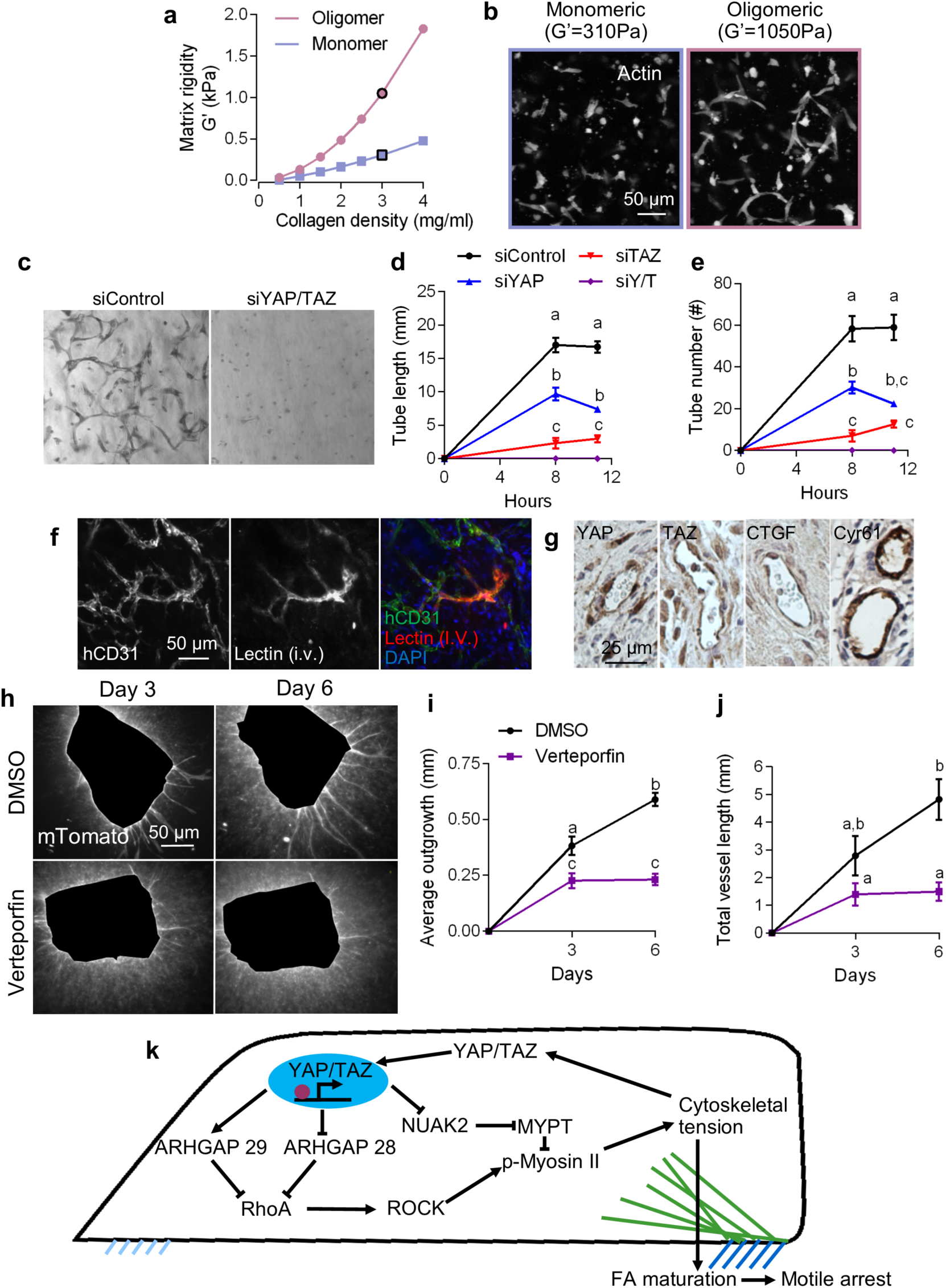
YAP and TAZ mediate endothelial tubulogenesis, vacuolation, and sprouting angiogenesis. ECFCs were plated on either matrigel for 12 hours or embedded in collagen matrices for 8 hours. **a,b,** Collagen matrix rigidity was modulated independent of density using either monomeric or oligomeric collagen,. **b,** Representative images of ECFC vasculogenesis in monomeric (G’ = 310 Pa) and oligomeric (G’ = 1050 Pa) matrices visualized with 488-conjugated phalloidin. **c,** ECFC vasculogenesis in oligomeric matrices (G’ = 132 Pa). **d,e**, Quantification of ECFC tubulogenesis on Matrigel at 8 and 12 hours (**d)** tube length (p < 0.0032) and number (p < 0.0003). n = 12, two-way ANOVA with Tukey’s *post hoc* test. **f,g,** ECFC embedded oligomeric matrices recovered from NOD-scid mice where used for immunohistochemistry. **f,** Functioning vasculature and human endothelium was visualized using Rhodamine-conjugated lectin and Alexa Fluor 647-conjugated anti-human CD31 antibody, respectively. **g,** Chromogenic HRP substrate DAB (brown) was used to visualize YAP, TAZ, CTGF, and CYR61 and counterstained with hematoxylin. **h,i,j,** Whole aortas were extracted from 5 CD57BL/6 with the ROSA 26-mTmG transgene, segmented, and embedded in oligomeric collagen for the aortic sprout assay. **h,** Representative images of mTomato expressing cell outgrowth from aortic explants at day 3 and 6. **i,** Average cell outgrowth measured as the average distance from the edge of the explant. n = 17-21 rings, p < 0.001, two-way ANOVA with Tukey’s *post hoc* test. **j,** The sum of all vascular sprout lengths from individual explants. p < 0.03, two-way ANOVA with Tukey’s *post hoc* test. Repeated significance indicator letters signify p > 0.05, while groups with distinct indicators signify p < 0.05. Summary statistics are represented as mean ± s.e.m. **k,** Schematic of the transcriptional negative feedback loop that regulates intracellular tension. Cytoskeletal tension is increased by ROCK activation of myosin II causing YAP and TAZ nuclear localization. Active YAP and TAZ inhibit ROCK-mediated MLC phosphorylation by transcriptional control of NUAK2 and ARHGAP 28, 29 expression, preventing overactivation of myosin II, enabling dynamic control of cytoskeletal tension.

ECFCs have potential as an autologous or allogeneic cell source for vasculogenic therapies^14,15^. To evaluate YAP/TAZ activation during transplanted ECFC vasculogenesis *in vivo*, we implanted human ECFC-laden collagen matrices in the subcutaneous space of NOD-scid mice. Establishment of a functional human neovascular plexus that inosculated with the host vasculature was demonstrated by confocal reconstruction of human CD31+ vessels that co-stained with intravenously-perfused Rhodamine-labeled UEA-I lectin (Figure 10f). Immunostained human neovasculature revealed nuclear YAP/TAZ and robust expression of target genes CTGF and Cyr61 *in vivo* (Figure 10g).

Finally, we evaluated the transcriptional role of YAP and TAZ in angiogenic sprouting *ex vivo* using verteporfin, a selective inhibitor of the YAP/TAZ-TEAD transcriptional complex^42^. We first demonstrated that 2 µM verteporfin treatment reproduced both the cytoskeletal and focal adhesion defects caused by RNAi depletion of YAP/TAZ (Supplementary Figure 9). Like YAP/TAZ siRNA, verteporfin treatment had no effect on total actin (p > 0.78), but increased actin polymerization (p < 0.0001) as well as vinculin (p < 0.0003) and paxillin incorporation (p < 0.02) into focal adhesions (Supplementary Figure 9).

To evaluate YAP/TAZ function during sprouting angiogenesis *ex vivo*, we quantified vessel outgrowth in the aortic sprout assay in monomeric and oligomeric collagen fibril matrices with shear moduli of 51 and 132 Pa, respectively. Aortic outgrowth and angiogenic sprouting were enhanced in stiffer oligomeric collagen matrices, but we did not observe a measurable difference in YAP/TAZ subcellular localization in response to 3D matrix rigidity (Supplementary Figure 10). Verteporfin treatment of vessel explants in oligomeric matrices significantly reduced non-vessel cellular outgrowth, a measure of three-dimensional cell migration (Figure 10h, i) and reduced endothelial neovessel length (Figure 10h, j), completely abrogating both cell migration and angiogenesis after day 3.

## Discussion

Whether existing proteins are sufficient to maintain migration, independent of transcription, is a matter of debate^43,44^. Here, we show that *de novo* gene transcription is essential for persistent ECFC motility and identify the transcriptional co-activators YAP and TAZ as regulators of migration through cell-intrinsic feedback control of Rho-ROCK-myosin II activity, partially through transcriptional repression of the MLCP regulator NUAK2 (Figure 10k). We validated the role of the Rho-ROCK-YAP/TAZ-NUAK2 signaling axis using RNAi and pharmacological inhibitors. Depletion or inhibition of MLCP regulatory kinase NUAK2 as well as myosin II/ROCK inhibition partially restored cell motility by relieving cytoskeletal tension. Notably, YAP/TAZ depletion, YAP/TAZ-TEAD inhibition, and global transcriptional/translational inhibition consistently increased stress fiber and focal adhesion maturation and arrested cell motility. Together, these data demonstrate that cytoskeletal dynamics, initiated by cell migration, activate YAP and TAZ to drive a transcriptional regulation program that feeds back to modulate cell mechanics, maintain a responsive cytoskeletal equilibrium, and prevent migration arrest.

Cells require new gene expression to replace consumed or degraded proteins. However, we found that transcription inhibition causes eventual motility arrest, not due to depletion of the components of the molecular clutch, but rather through dysregulated cytoskeletal polymerization and focal adhesion formation and maturation. This is consistent with the reported stability of the molecular clutch proteins. For example, vinculin and talin have a half-life of 18-21 hours^45^, β-actin has a half-life of 48 hours^46^, while myosin contractile motors in muscle are stable for days^47^. Consistently, we observed stress fiber and focal adhesion formation even after long term transcription inhibition. Further, myosin tension generation and subsequent focal adhesion reinforcement occurs more rapidly than *de novo* protein synthesis^2,23^. However, there is ample evidence that immediate early genes are transcriptionally upregulated during migration and enhance cell motility^48–50^. These observations led us to explore the transcriptional mechanisms by which migration initiation modulates cytoskeletal remodeling to enable persistent motility.

We found that YAP/TAZ-mediated transcription is essential to moderate cytoskeletal mechanics, and YAP/TAZ depletion or inhibition mimics the effects of global transcription inhibition. Thus, we conclude that YAP and TAZ act to dissipate cytoskeletal tension in part through a cell-intrinsic feedback mechanism dependent on NUAK2 control of myosin II activation. These data contribute to an emerging but conflicting literature regarding the feedback functions of YAP and/or TAZ in regulating cellular and tissue tension. Some recent reports implicate YAP in feed-forward promotion of cytoskeletal tension^34,51^, potentially through outside-in feedback through regulation of extracellular matrix production and subsequent mechanosensation^52–54^. In contrast, our data are consistent with evidence from cells that do not produce extensive extracellular matrix, implicating YAP/TAZ-mediated ARHGAP expression in suppression of actin polymerization and cytoskeletal tension^36^. While it is clear that the functions and relative roles of YAP and/or TAZ are highly cell type- and context-dependent^55^, synthesizing the available data, we conclude that YAP/TAZ mediate both cell-autonomous and outside-in feedback systems to modulate cell and tissue tension.

Here, we identify a cell-autonomous feedback pathway by which YAP/TAZ suppress cytoskeletal tension through Rho-ROCK-myosin II signaling, verified through orthogonal mechanical and biochemical measurements. We further validated this negative feedback system through both loss-of-function and rescue experiments featuring pharmacologic and RNAi-based approaches, and identify the YAP/TAZ-TEAD-dependent target gene, NUAK2, as a novel negative regulator of cell migration through cytoskeletal tension in ECFCs. These data do not suggest that cytoskeletal tension inhibits migration, per se, but rather that transcriptional programs modulate actomyosin activity. We identify YAP/TAZ-mediated repression of NUAK2 as a key modulator of cytoskeletal tension. We also observe simultaneous upregulation and downregulation of ARHGAP28 and 29, respectively. These data both conflict and conform to previously published literature^34,36^, but support the common hypothesis that YAP and TAZ can act both as co-activators and co-repressors of gene expression, depending on cell type and environment^56–58^. While YAP/TAZ depletion largely recapitulates the effects of global transcription inhibition, other mechanotransductive factors may also contribute to parallel or interacting feedback loops. For example, the transcriptional co-activator MRTF transactivates SRF-dependent gene expression, including actomyosin genes, MLC2 and β-actin, and focal adhesion components talin, vinculin, and zyxin^59,60^. Similarly, AP-1 transcribes cytoskeletal regulators gelsolin like capping protein (CapG) and Kelch related protein 1 (Krp1) while downregulating fibronectin^61^. Interestingly, there is significant overlap between MRTF, AP1, and TEAD occupancy at inducible genes^62,63^. Thus, transcriptional mechanotransductive mechanisms are interdependent and further study will be required to clarify how complex multi-transcription factor dynamics tune the cytoskeleton and regulate both motility and mechanosensation.

YAP and TAZ are essential during development and can have either convergent or divergent functions depending on the experimental context. Global deletion of YAP is embryonic lethal, due to impaired vasculogenesis, whereas TAZ knockout mice survive birth but only 50-65% reach adulthood due to polycystic kidney disease^64^. YAP is the most well-studied of the pair, and was hypothesized to have the more potent effect on transcriptional activation, due primarily to the presence of a PXXΦP motif^65^. However, a recent structural analysis found a distinct TAZ-TEAD conformation that confers similar or greater transcriptional activity compared to YAP-TEAD^65^. *In vivo* evidence from our group and others have found that YAP vs. TAZ functional redundancy is cell-type specific. For example, YAP is essential, but TAZ dispensable, during cardiac development^66^, while YAP and TAZ are compensatory in bone development with TAZ exhibiting greater potency^55^. Interestingly, we observe a greater effect of TAZ ablation on cytoskeletal feedback, cell migration, and vessel formation in ECFCs. These data are consistent with recent descriptions of the roles of YAP and TAZ in endothelial cell migration and vascular integrity^67^. However, others have observed an increase in cell migration when YAP was depleted by shRNA in HUVECs and implicate a cytosolic YAP-CDC42 signaling axis, though the mechanism^68^ remains unclear. While further research will be necessary to dissect potentially distinct co-effector interactions or transcriptional targets of YAP and TAZ, these data contribute to the emerging evidence of crosstalk between transcriptional activity and cytoskeletal dynamics.

This study also provides new mechanistic understanding to explain recent observations regarding the roles of YAP and TAZ in cardiovascular function *in vivo*. These studies show that endothelial-specific YAP/TAZ deletion is embryonic lethal^69,70^, with defects in retinal angiogenesis, liver vascularization, and hindbrain hemorrhage^68^. Prior data implicates defects in proliferation^35^, tip cell sprouting^68,70,71^, and vascular integrity^69^. Our findings establish a new YAP/TAZ-Rho-ROCK-myosin II feedback axis as a critical mechanism for neovascular function and points to transcriptional-cytoskeletal feedback as a key regulator of cell motility.

## Methods

### Cell culture and transfection

ECFCs were isolated from umbilical cord blood as previously described^14,15^ and cultured in endothelial growth medium (EGM-2 with bullet kit; Lonza, CC-3162) supplemented with 1% penicillin/streptomycin (Corning) and 10% defined fetal bovine serum (Thermofisher), referred to as full medium. Cells were seeded on collagen (5 µg/cm^2^) coated tissue culture polystyrene (TCPS) and maintained at 37° Celsius and 5% CO_2_. ECFCs were released from culture dishes using TrypLE^TM^ Express (Gibco) and used between passages 6 and 8.

ECFCs were depleted of YAP and TAZ using siRNA loaded lipofectamine RNAimax (Invitrogen) according to the manufacturer’s instructions. Briefly, ECFCs were seeded on collagen coated 6 well-plates, 10^5^ cells per well, in antibiotic free medium and kept in culture for 24 hours followed by transfection at approximately 50% confluence. Transfection was carried out using a final concentration 0.3% (v/v) lipofectamine RNAimax with 15 pmol RNAi duplexes (custom oligonucleotides; Dharmacon) per well. Transfected ECFCs were used 24-48 hours post-transfection.

ON-TARGET plus non-targeting siRNA and SMARTpool NUAK2 siRNA were obtained from Dharamacon. Custom siRNA were created based on sequences previously described^11^: YAP1, sense, 5’-GACAUCUUCUGGUCAGAGA-3’, YAP1, anti-sense, 5’-UCUCUGACCAGAAGAUGUC -3’; YAP2, sense, 5’-CUGGUCAGAGAUACUUCUU -3’, YAP2, anti-sense, 5’-AAGAAGUAUCUCUGACCAG -3’; TAZ1, sense, 5’-ACGUUGACUUAGGAACUUU -3’, TAZ1, anti-sense, 5’-AAAGUUCCUAAGUCAACGU -3’; TAZ2, sense, 5’-AGGUACUUCCUCAAUCACA -3’, TAZ2, anti-sense, 5’-UGUGAUUGAGGA AGUACCU -3’.

### Collagen synthesis and characterization

Collagen oligomers and monomers were synthesized and polymerized as previously described^16,17,72^. Briefly, porcine skin collagen was isolated using acetic acid extraction yielding a viscous collagen composition containing both oligomeric and monomeric collagen^73^. Monomeric collagen was purified by salt precipitation, selectively removing oligomeric collagen from the solution^17^.

Collagen mechanical properties were defined as previously described^72^. Storage and loss moduli were measured in oscillatory and shear and compression on a stress controlled AR2000 Rheometer (TA instruments). Collagen samples were polymerized *in situ* at 37° Celsius for 30 minutes and then tested with a shear strain sweep from .01% to 5% at 1 Hz. Following the strain sweep compressive modulus (E_c_) was measured in unconfined compression at a strain rate 20 µm/s (2.76% strain per second). Stress was calculated as the normal force divided by plate area (12.57 cm^2^). E_c_ was then calculated as the slope of the stress strain curve.

### Animal Handling

All animal experiments were approved by the institutional animal care and use committee (IACUC) at the University of Notre Dame and Indiana University School of Medicine.

10^6^ cord blood derived ECFCs were re-suspended in 250 ul collagen gel (G’ = 200Pa, Geniphys, Standarized Oligomer Polymerization Kit) plus 10% human platelet lysate (Cook) on ice and then polymerized for 30 minutes at 37° C in a well of 48 well plate. Matrices were covered with 500ul culture medium until transplantation. Cellularized matrices were transplanted into the abdominal flanks of 6-12 week old NOD/SCID immunodeficient mice anesthetized by inhaled isoflurane under aseptic conditions. After 14 days, 100 ul of Rhodamine labeled Ulex Europaeus Agglutinin I (UEA I referred to in this report as Lectin, Vector Laboratories) were intravenously injected into the transplanted mice 30 minutes before the mice were euthanized. The grafts were collected from the mice and fixed in 4% paraformaldehyde at 4° C overnight and prepared for immunofluorescence or paraffin embedding.

Whole mouse aortas were extracted as previously described^74^. Briefly, 4-6 week-old C57BL/6 mice with or without the mTomato/mGFP (mTmG) transgene were anesthetized by inhalation of 5% isoflurane in oxygen followed by physical euthanasia by bilateral thoracotomy. Whole aortas from the aortic arch to the abdominal insertion were extracted and cleaned of fat and branching vessels then flushed with dPBS containing 10 U/mL of heparin sodium (Hospira). Aortas were sectioned into 0.5 mm rings and serum starved overnight in EBM-2 with 1% penicillin/streptomycin. Aortic rings were encapsulated in oligomeric collagen (G’ = 132 Pa) in 96 well plates with full medium containing either verteporfin (Sigma) or an equal volume of dimethyl sulfoxide (DMSO, Sigma). Fluorescent z-stacks of aortic rings expressing mTmG were taken with a Leica DMi8 0, 3, and 6 days after polymerization. Sprouting aortas not expressing mTomato were fixed with 4% paraformaldehyde for immunofluorescence.

### Polyacrylamide hydrogels

Polyacrylamide hydrogels were prepared as previously described^75^ with slight modifications. Briefly, 24×50 mm #1 glass coverslips were washed with soap and water and rinsed in ethanol. Coverslips were functionalized using 0.5% (v/v) 3-(Trimethoxysilyl)propyl methacrylate (sigma) in ethanol. Coverslips were cut to 24×20 mm sections to fit in 6 well plates. Polyacrylamide precursor solutions were prepared from 40% acrylamide (EMD Millipore), 2% Bis-acrylamide (amresco), Tetramethylethylenediamine (TEMED; ThermoFisher), and ammonium persulfate (APS; ameresco). 45 µL of polyacrylamide precursor solution was pipetted on to hydrophobic glass slide and topped with the functionalized glass coverslips and allowed to polymerize for 30 minutes.

Hydrogels were functionalized with extracellular matrix (ECM) using techniques described previously^75^. Hydrogels were first treated with hydrazine hydrate (sigma) overnight then washed with deionized water followed by 1 hour in 5% (v/v) acetic acid then 1 hour in deionized water. Collagen (MP Biomedicals) in 50mM sodium acetate buffer (pH 4.5; Sigma) with 4 mg/mL sodium (meta)periodate (Sigma) for half an hour in the dark. ECM was then applied to the hydrazine hydrate functionalized hydrogels for 1 hour. Hydrogels were thoroughly washed in deionized water and equilibrated in PBS overnight. Hydrogels were sterilized for 15 minutes under a germicidal UV lamp then washed in EBM-2 and equilibrated in full medium for at least 8 hours. For immunofluorescence and migration experiments 8.5 × 10^3^ cells per cm^2^ were seeded per hydrogel.

### Migration assays

Migration assays were performed on confluent layers of transfected and/or inhibitor treated cells. 24 hours post-transfection cells were washed twice in endothelial basal medium (EBM-2) then serum-starved in EBM-2 for 2 hours. The collective migration wounding assay was performed as described previously^76^. Migration was initiated by scratching monolayers were vertically and horizontally with the tip of a 200 µL pipet tip followed by two washes in EBM-2 and addition of full medium. Actinomycin D (Sigma), puromycin (Sigma), WZ4003 (MedChem Express), Y-27632 (Tocris), and blebbistatin (Sigma) were added into basal medium 1 hour after starting serum starvation and in full medium after initiation of migration. Verteporfin (Sigma) treatment in conjunction with serum starvation resulted in significant cell death, to prevent this cells treated with verteporfin were not serum starved. ECFCs were treated with mitomycin C (Tocris) diluted in basal media during the serum starve, prior to migration. Phase images of migration were taken on a Leica DMi8 or Zeiss Axio Observer Z1. After 8-12 hours cells were fixed for immunofluorescence.

Live migration was performed on ECFCs expressing mTomato or EGFP in confluent or sparse conditions on collagen coated 35mm dishes. GFP-expressing ECFCs were transfected with non-targeting control siRNA, whereas mTomato-expressing ECFCs were transfected with siRNA targeting YAP and TAZ. Cells were seeded in a polydimethylsiloxane stencil with 3×5 mm channels separated by 0.75 mm gaps. 2 hours after plating the barrier was released and seeded regions where imaged in 15 minutes intervals for 10 hours using a Zeiss Axio Observer Z1 inverted microscope with an automated stage. Cells were maintained in an incubation chamber at 37º Celsius, 5% CO_2_, and 95% relative humidity for the duration of the experiment.

### Immunofluorescence

Cells were washed twice in EBM-2 and fixed in 4% paraformaldehyde (Alfa Aesar) diluted in cytoskeletal stabilization buffer containing 10 mM 2-(N-morpholino) ethanesulfonic acid, 150 mM potassium chloride, 3 mM magnesium chloride, 2 mM ethylene glycol-bis(β-aminoethyl ether)-N,N,N’,N’-tetraacetic acid, and .3 M sucrose for 20 minutes at room temperature. Cells were permeabilized and blocked in PBS containing 0.03% triton x-100 (amresco) with 5% goat serum (Cell Signaling) for one hour. Isolation of stable focal adhesions *in situ* was done by adding 0.05% triton x-100 to the cytoskeletal stabilization buffer to remove the soluble and weakly adhered fraction of focal adhesions, as described previously^39^.

Fixed samples were incubated with antibodies diluted in PBS with 1% BSA: monoclonal YAP ab (1:200, Cell Signaling, 14074), polyclonal TAZ ab (1:250, Cell Signaling, 4883), monoclonal Vinculin ab (1:200, Cell signaling, 13901), MLC (1:200, Cell Signaling, 3672), pMLC (1:200, Cell signaling, 3675), α-tubulin (1:2000, Cell Signaling, DM18), GM130 (1:200, Cell Signaling, D6B1), Rab7 (1:100, Cell Signaling, D95F2), FAK (1:100, Cell Signaling, 3285), pFAK (1:100, Cell signaling, D20B1), paxillin (4 µg/mL, abcam, 80578), polyclonal Alexafluor 594 conjugated anti-rabbit IgG (1:400, Cell Signaling, 8889), and polyclonal Alexafluor 488 conjugated anti-mouse IgG (1:400, Cell Signaling, 4408). F-actin was stained using Alexa fluor 488 conjugated phalloidin (1 unit/mL; Life Technologies) and G-actin stained with Alexa fluor 594 conjugated DNase I (0.3 µM; Life Technologies) for 15 minutes. Epifluorescence images of fixed samples were taken on either a Leica DMi8 or Zeiss Axio Observer.

### Integrin endocytosis assay

β1-integrin endocytosis was performed as previously described^5^ Integrin internalization was performed on ECFCs sparsely plated on collagen coated glass cover slips. 24 hours after transfection cover slips were inverted on to 50 µL full medium droplets with β1-integrin ab (10 µg/mL, abcam, 12G10) on UV sterilized parafilm. Coverslips were then placed a 4° C for 45 minutes to allow β1-integrin ab to target active surface integrins while preventing active integrin endocytosis. Coverslips were returned to 6 well plates with pre-warmed full medium and washed 3 times with full medium at 37° C for 30 minutes to allow internalization of integrin-ab conjugates. Coverslips were washed 3 times in 4° C EBM-2 (pH 4.0) to denature any remaining surface-bound antibodies. Cells were briefly washed with physiologic pH EBM-2 (pH 7.4) twice then fixed as previously described in cytoskeletal stabilization buffer. Antibody detection and additional immunostaining was performed as described in the previous section.

### Single-cell nanoindentation

ECFC stiffness at different distances from the leading edge of migrating cells was tested using a PIUMA CHIARO nanoindenter system (Optics11)^77^. A colloidal probe cantilever with a tip radius and spring constant of 9.5 µm and 0.068 N/m, respectively, with a loading velocity of 2 µm/s was used in this study. Before testing, the sensitivity calibration of the cantilever was conducted by indenting a hard surface (i.e. a petri dish). Briefly, cell stiffness at the leading edge of the wound and in 100 µm bins from the leading edge were tested. Two control and three YAP/TAZ depleted samples were tested with a total indentation of 15-32 cells at each measuring location. A customized MATLAB code (The MathWorks, Inc.)^77^ was developed to determine contact points between the probe and cells and to identify Young’s moduli of the cells using the Hertz contact model:

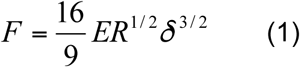

where *F* is applied force, *δ* is indentation depth, *R* is the radius of the colloidal probe, and *E* is Young’s modulus of the cells. The cells were assumed to be incompressible (i.e. Poisson’s ratio of 0.5).

### Immunohistochemistry

Collagen matrices containing lectin labelled endothelial cell were stained with 1:100 Alexa Fluor 647 conjugated mouse anti human CD31 antibody (BD pharmigen clone WM59) at 4 °C overnight. Next, the samples were cut into 0.3-0.5mm thick pieces and mounted onto Superfrost Plus Gold microscope glass slides (Thermo Fisher Scientific) with ProLong® Gold Antifade solution with DAPI (Thermo Fisher Scientific). Fluorescent pictures of vessels were taken on an Olympus II confocal microscope.

Paraffin embedded tissues were sectioned, deparaffinized and rehydrated. Heat-induced epitope retrieval was performed by incubating sections in sub-boiling 10 mM citrate buffer (pH 6.0) for 10 minutes followed by washes in deionized water. Non-specific binding was blocked using horse serum from the Vectastain elite avidin-biotin conjugation kit (Vector, ABC kit) and endogenous peroxidase activity was quenched using 0.3% hydrogen peroxide in deionized water. Sections were incubated overnight 4 °C with antibodies targeting YAP (1:400), TAZ (1:400), CTGF (1:400), or Cyr61 (1:400). The ABC kit universal secondary and biotinylated horseradish peroxidase containing reagents were added according to the manufacturer’s instructions. Antibody conjugation was detected using ImmPACT DAB peroxidase substrate. Sections were counterstained with hematoxylin and eosin (Sigma), coversliped, and imaged with a Nikon 90i.

### Western Blot

Cells were washed in ice cold dPBS then lysed in 2x lammeli buffer (Alfa Aeasar) 24-30 hours post-transfection. Lysate was denatured by boiling samples for 5 minutes followed by centrifugation at 12,000xg for 15 minutes. Lysate was separated based on molecular weight (MW) using electrophoresis on precast 4-12% Clearpage SDS gels (CBS Scientific) in conjunction with a low range molecular weight ladder (amresco) in TEO-tricine running buffer (CBS Scientific). Proteins were transferred to PVDF membranes (amresco) in tris-glycine transfer buffer (CBS Scientific). Membranes were washed in TBST (GeneTex), blocked in 5% BSA in TBST for one hour, and incubated overnight at 4° Celsius with primary antibodies targeting YAP (1:250), TAZ (1:1000), or GAPDH (1:2000) diluted in blocking buffer. The following day membranes were washed in TBST followed by incubation with HRP-conjugated secondary (1:3000, Cell Signaling, 7074) for one hour at room temperature, washed in TBST, and detected with chemiluminescent substrate (Pierce). Images were taken on Chemi-Doc It®^2^ (UVP) and densitometry of bands performed on UVP Chemi-Doc It® software, semi-quantitative comparisons were made after normalizing to GAPDH. Membranes were stripped using a mild stripping buffer pH 2.2 (1.5% glycine (amresco), 1% tween-20, 0.1% SDS (amresco)) twice followed by repeated washes in PBS and TBST.

### RT-qPCR

Total RNA was isolated and purified using the RNeasy mini kit (qiagen). 0.5 µg of total RNA was reversed transcribed using the TaqMan reverse transcription kit (Life Technologies) using the manufacturer’s instructions in a thermal cycler eco (Eppendorf). cDNA was mixed with iTaq Universal SYBR supermix (Biorad) and 0.4 µM forward and reverse primers in wells of 96 well PCR plate (Biorad). followed by amplification and quantification with in a CFX connect real time PCR system (Biorad).

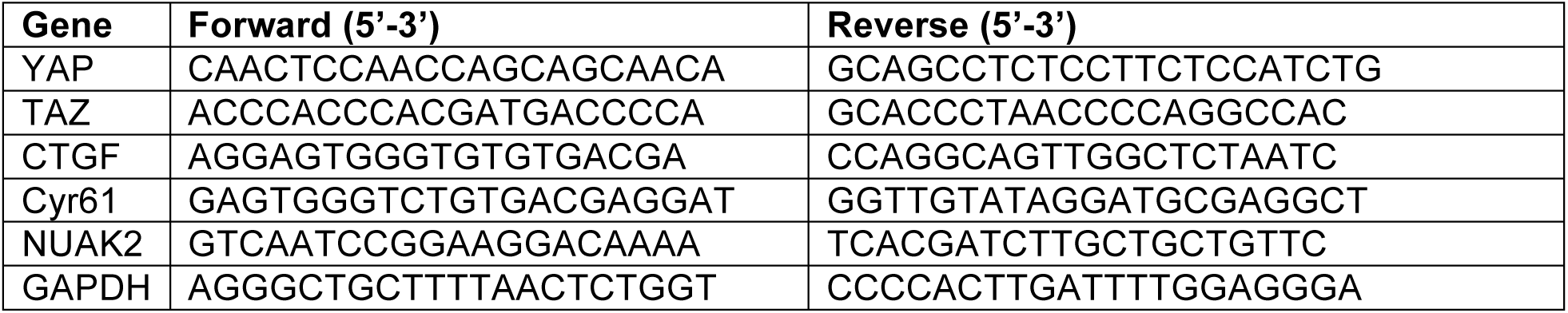

### Image analysis

All image analysis (morphometrics, intensity measurements, and individual cell tracking) were performed using an open access NIH software platform, FIJI^78^. Live migration tracking was performed using a semi-automated tracking algorithm, track mate^79^. Cells within 100 µm of the leading cell were considered front cells, all other cells were considered trailing.

Golgi polarization relative to wound edge was found by finding the angle between the horizontal direction of migration and the position vector between the centroid of nuclei and the associated Golgi in matlab using positional information obtained in image J. Leading edge cells where defined as cells with 50 µm of the front most cell.

YAP and TAZ activation as a function of cell density was performed on images of cells up to 1.2mm from the actively migrating front. Images were binned into adjacent 100 µm × 950 µm ROI’s starting at the leading cell. Total and activated YAP or TAZ is the total integrated fluorescent intensity in an ROI with or without the nuclear fraction digitally removed. Cell area in an ROI is the average area of 5 cells per ROI and density is the number of DAPI stained nuclei across the ROI.

Fluorescent intensity of immunocytochemistry was measured in epiflourescent images. Single cell fluorescence was measured in cells with minimal cell contact, at the leading edge, whereas focal adhesion morphometrics were performed in sparsely plated cells. Florescent intensity measurements were taken from cells across at least two independent experiments, each independent experiment was performed in duplicate or triplicate, measurements were taken from each sample and normalized by the average intensity of the control for the experiment.

### Statistics

All statistical analyses were performed on Graphpad Prism 6 statistical analysis package. Data is presented with data points were possible and mean ± standard error unless indicated otherwise. Multiple comparisons were made using analysis of variance (ANOVA) with Tukey’s *post hoc* test for pairwise comparisons of normally distributed homoscedastic data. Data that did not meet the ANOVA criteria were analyzed by Kruskal-Wallis with Dunn’s *post hoc* test. Letters denote significant differences between groups, p-values < 0.05 were considered significant and exact p-values for the comparison with highest significant p-value is recorded in the figure legend.

### Data/code availability statement

The datasets and code in the current study are available from the corresponding author on reasonable request.

## Acknowledgements

We would like to thank William Chris Shelley for his technical assistance with ECFC culture and vasculogenesis assays. Lance Davidson for insightful discussions. Jason Burdick for providing cells.

Funding: This project was supported in part by American Heart Association Grant 16SDG31230034 (to J.D.B.), by U.S. National Institutes of Health (NIH) National Center for Advancing Translational Sciences Grant UL1TR001108 (to J.D.B.) and National Institute of Arthritis and Musculoskeletal and Skin Disesases Grant AR071559 (to J.D.B.), by the National Science Foundation Grant 1435467 (to J.D.B.), and the National Science Foundation’s Science and Technology Center for Engineering MechanoBiology, grant CMMI-1548571 (to J.D.B.). The authors declare no conflicts of interest.

## Author Contributions

D.M., J.D., D.T.N., and Y.L.: performed the experiments. D.M. and J.B. conceived and designed the experiments and wrote the manuscript. S.V-H., P.Z., and M.Y.: provided reagents, cells, instrumentation, and advice for completing experiments. J.D., D.T.N., Y.L., S.V-H., P.Z., and M.Y.: reviewed and provided feedback on the manuscript.

---

**Supplementary Figure 1:**
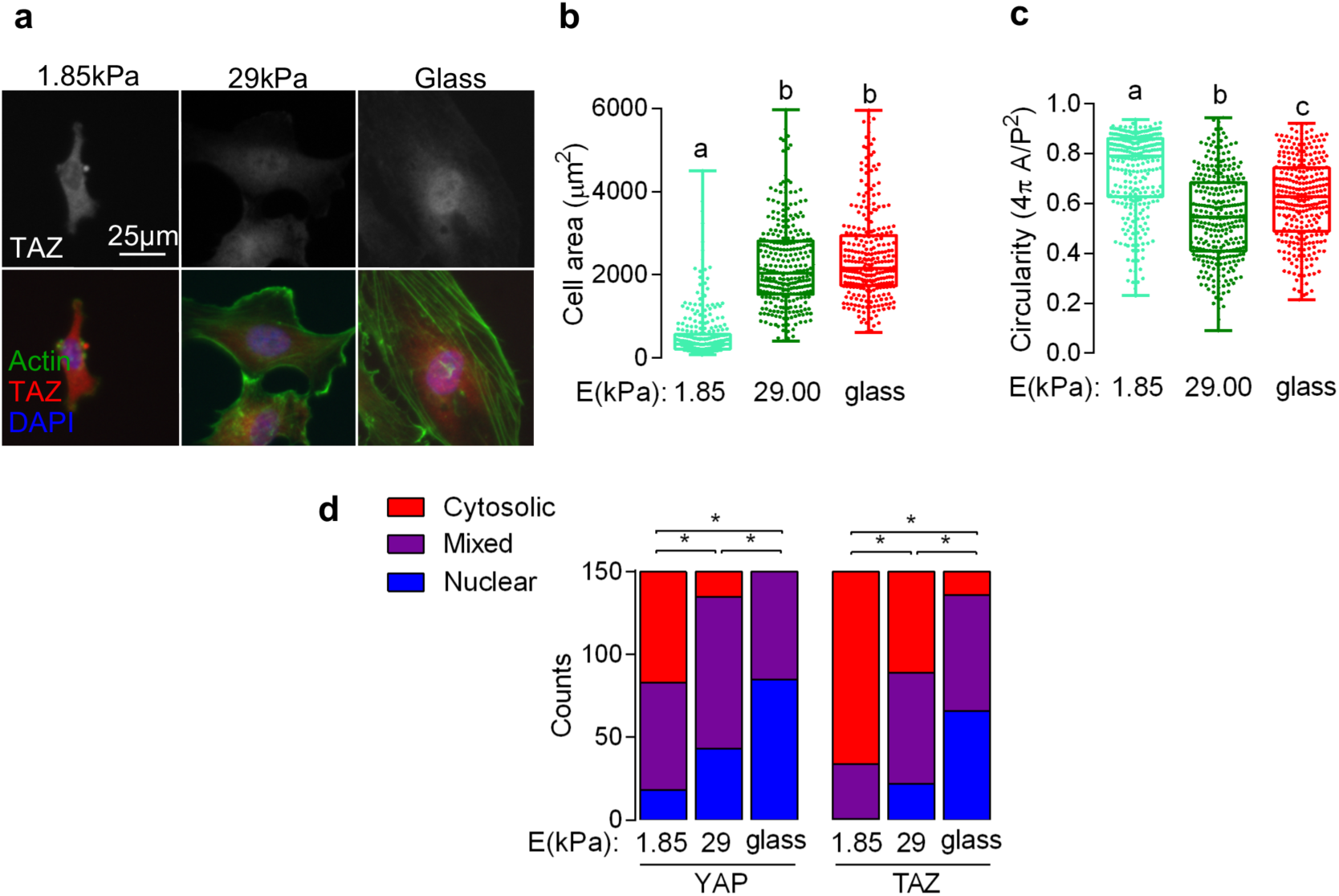
YAP and TAZ nuclear localization are sensitive to rigidity in ECFCs. ECFCs were seeded at 8.5 × 10^3^ cells per cm^2^ on collagen coated soft or stiff polyacrylamide or glass for 24 hours then fixed and stained for YAP (not shown) and TAZ. **a,** Representative images actin, TAZ, and nuclei visualized with Alexa Fluor 488- and 594-conjugated phalloidin and secondary and DAPI, respectively. **b,** Individual cell area and **c,** circularity as a function of substrate elastic modulus. **d,** Cumulative counts of YAP or TAZ subcellular localization based on qualitative assessment, n = 150, p < 0.0001, Chi-square test.

**Supplementary Figure 2:**
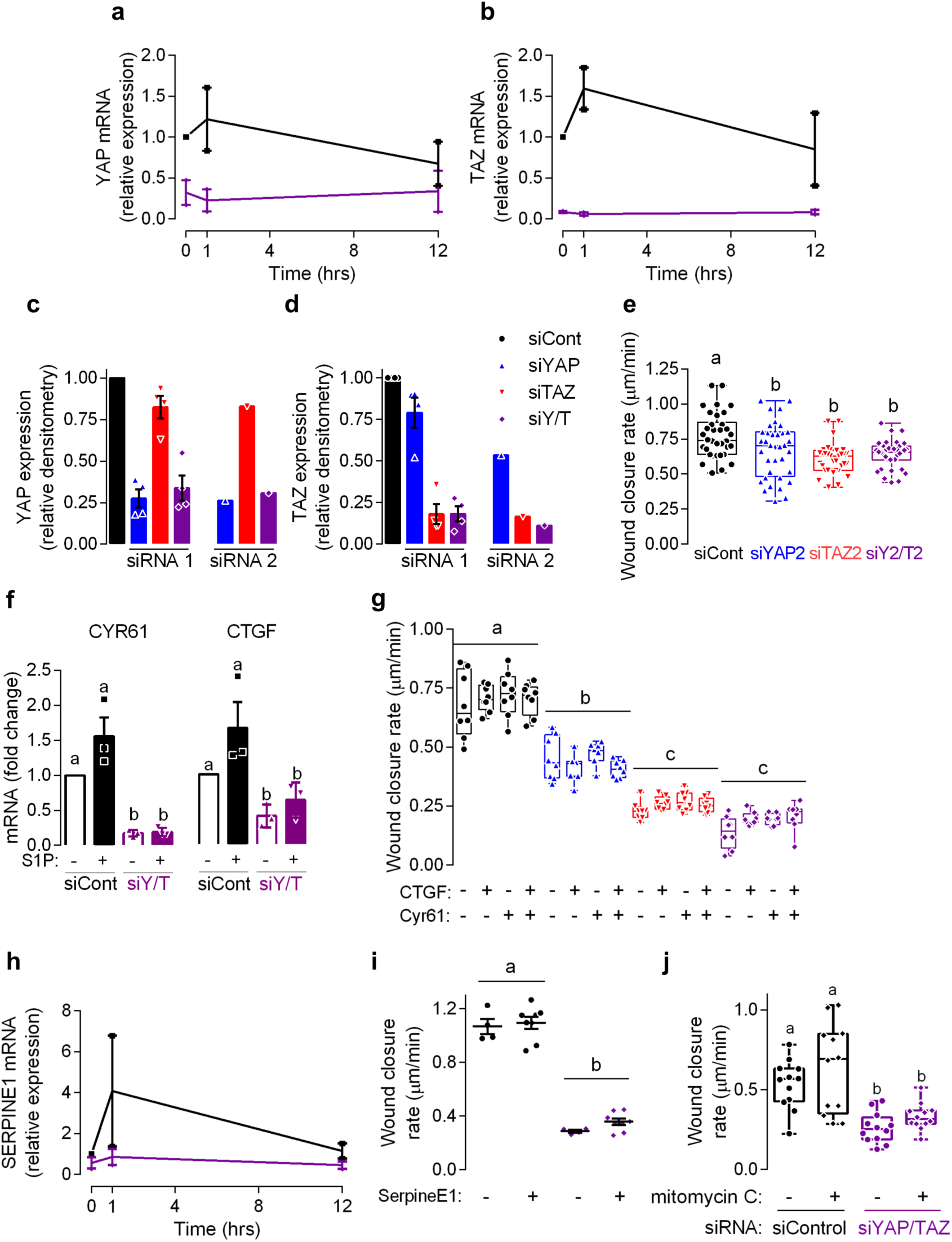
Reconstitution of YAP/TAZ-dependent angiocrines CTGF, Cyr61, or SERPINE1 does not rescue ECFC migration. Confluent ECFCs were scratched and lysate collected for gene expression 0, 1, and 12 hours after initiation of migration. **a,** YAP and **b,** TAZ mRNA expression in migrating ECFCs depleted of YAP and TAZ, n = 4. **c,** YAP and **d,** TAZ protein expression after depletion with siRNA #1 and #2. **e,** Wound closure rate of confluent ECFCs imaged over 12 hours after depletion of YAP and TAZ with siRNA #2, n = 34, p < 0.01, ANOVA with Tukey’s *post hoc* test. **f,** CTGF and Cyr61 gene expression after treatment with sphingosine-1-phosphate for 1 hour followed by lysis and RNA collection for RT-qPCR. n = 2 independent experiments, p < 0.01, ANOVA with Tukey’s *post hoc* test. **g,** Wound closure rate of cells treated with 50 and 100 ng/mL of CTGF and/or Cyr61. Groups treated with 50 or 100 ng/mL were combined for analysis, as there were no differences in wound closure between those treatments, n = 7−8, p < 0.0001, ANOVA with Tukey’s *post hoc* test. Migrating ECFCs were lysed at 0, 1, and 12 hours after the initiation of migration and lysate use for RT-qPCR. **h,** SERPINE1 gene expression during migration. N = 2 independent experiments. **i,** Wound closure rate after treatment with 50 or 100 ng/mL of SERPINE1. Groups treated with 50 or 100 ng/mL were combined for analysis, as there were no differences in wound closure between those treatments, n = 4−8, p < 0.0001, ANOVA with Tukey’s *post hoc* test. **j**, ECFCs were pre-treated with 20 µg/mL mitomycin C (mito C) during serum starvation, prior to initiation of cell migration. Wound closure rate (µm/min) was measured after 8 hours, n = 12, p < .02, ANOVA with Tukey’s *post hoc* test. Summary statistics in **a**, **b**, **c**, **d**, **g**, **f** and **h** are represented as mean ± s.e.m.

**Supplementary Figure 3:**
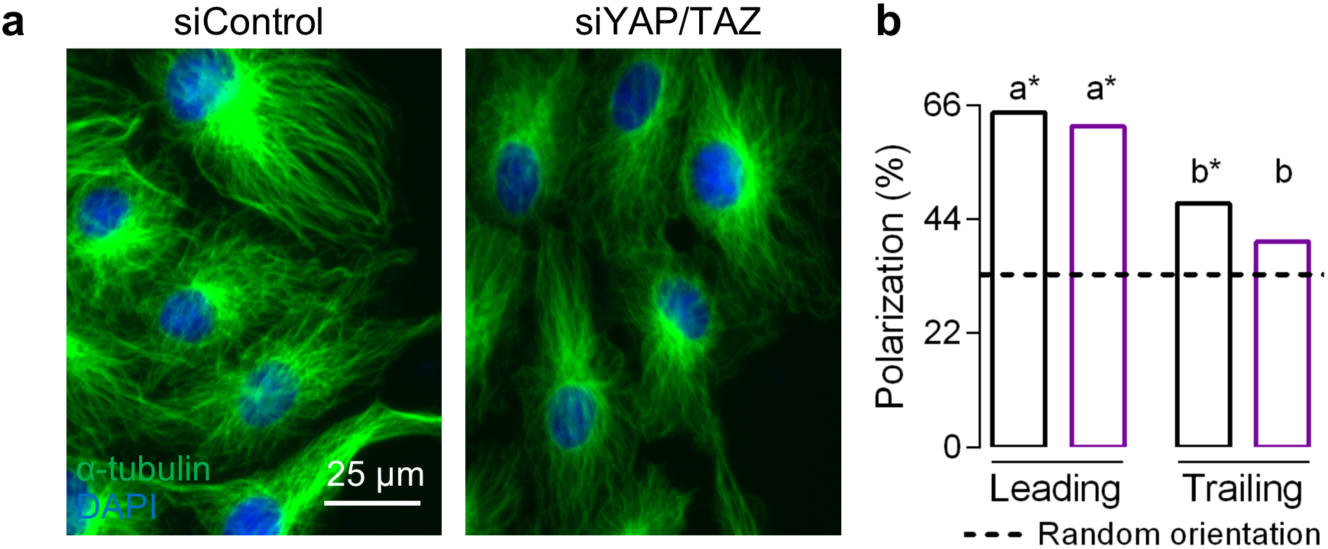
YAP and TAZ are important for collective polarization, but dispensable for single cell polarization. Confluent ECFCs were scratched and allowed to migrate for 8 hours followed by fixation for immunofluorescence and polarization analysis. **a,** α-tubulin and nuclei visualized by Alexa Fluor 488-conjugated secondary and DAPI, respectively. **b,** Percentage of cells with Golgi polarized in the direction of the wound edge in leading and trailing regions, n = 100-105 leading, 350-360 trailing, and 342-421 confluent cells, p < 0.02, * p < 0.002, ANOVA with Bonferroni *post hoc* test.

**Supplementary Figure 4:**
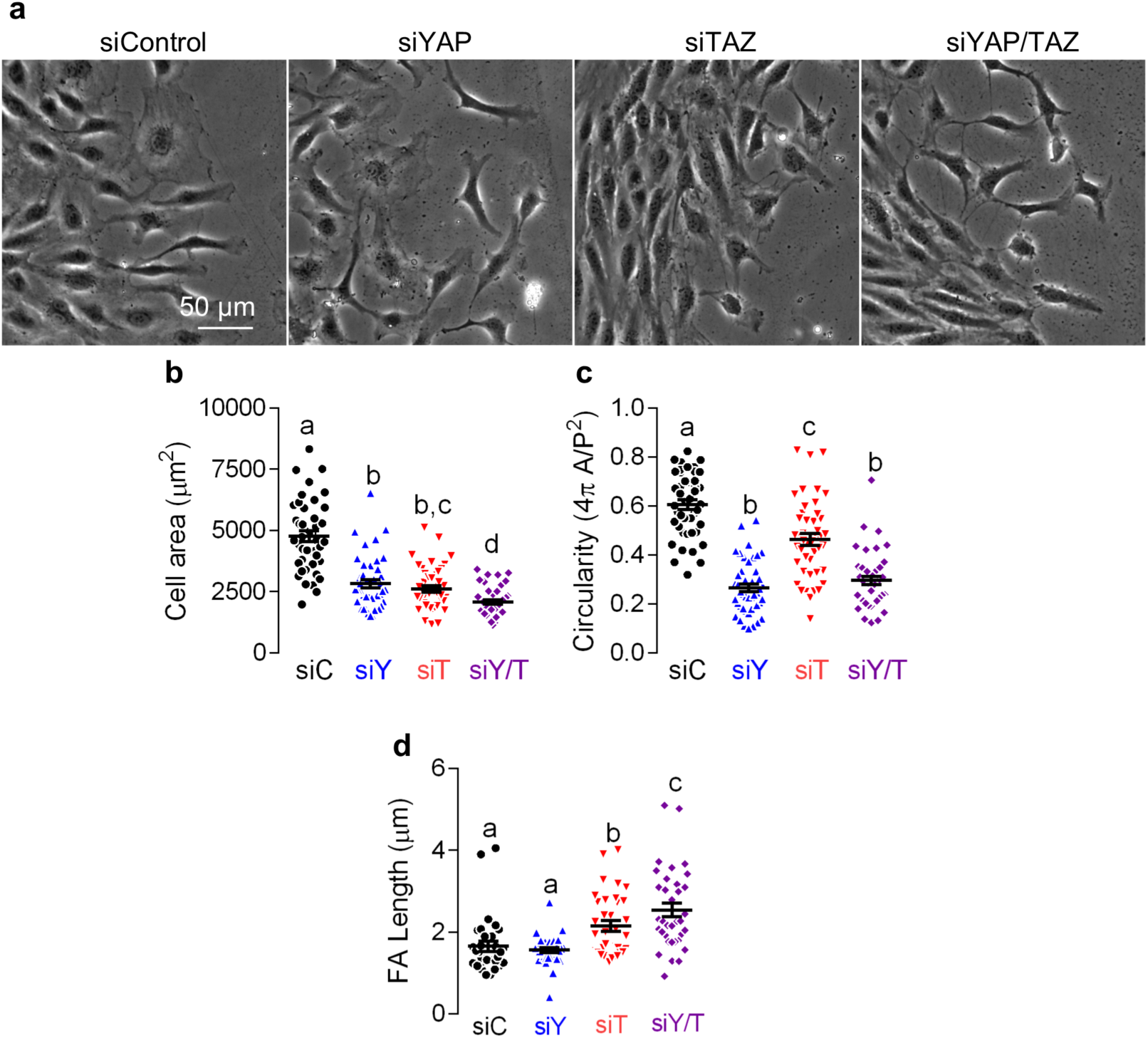
YAP and TAZ control cell morphology, in part through focal adhesion maturation. Confluent ECFCs were scratched and allowed to migrate for 8 hours then fixed for morphological analysis. **a,** Representative phase images of cell morphology. **b,** Cell area, p < 0.004, and **c,** circularity, p < 0.0001, n = 45, ANOVA with Tukey’s *post hoc* test. **d,** Average vinculin+ focal adhesion length from data described in Figure 6c,, n = 33-35, p < 0.007, ANOVA with Tukey’s *post hoc* test. Summary statistics in **b**, **c**, and **d** are represented as mean ± s.e.m.

**Supplementary Figure 5:**
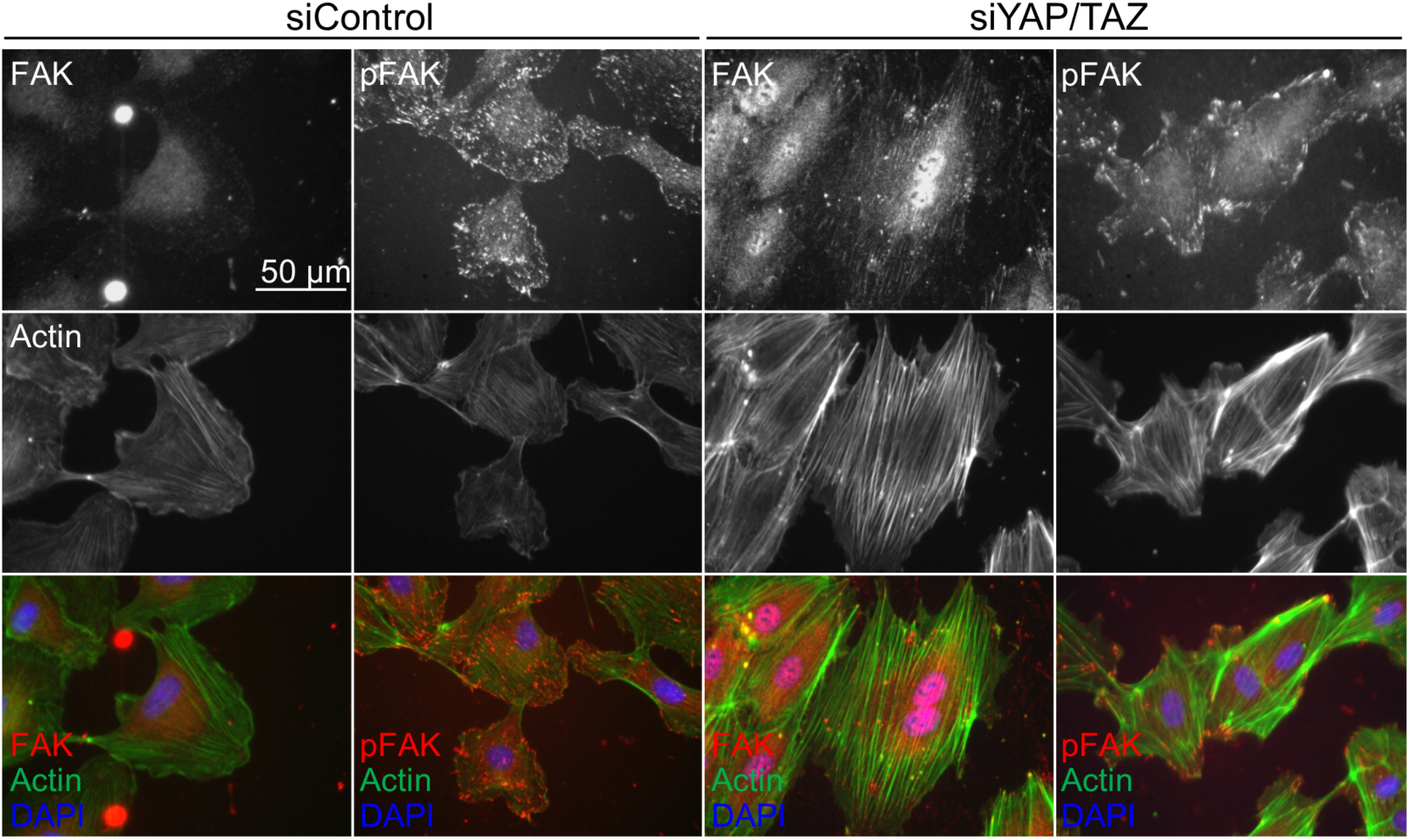
YAP/TAZ-mediated cytoskeletal changes drive changes in FAK and pFAK localization. Confluent ECFCs were scratched and allowed to migrate for 12 hours then fixed for immunofluorescence. FAK, pFAK, actin, and nuclei were visualized with Alexa Fluor 594-conjugated secondary, Alexa Fluor 488-conjugated phalloidin, and DAPI, respectively.

**Supplementary Figure 6:**
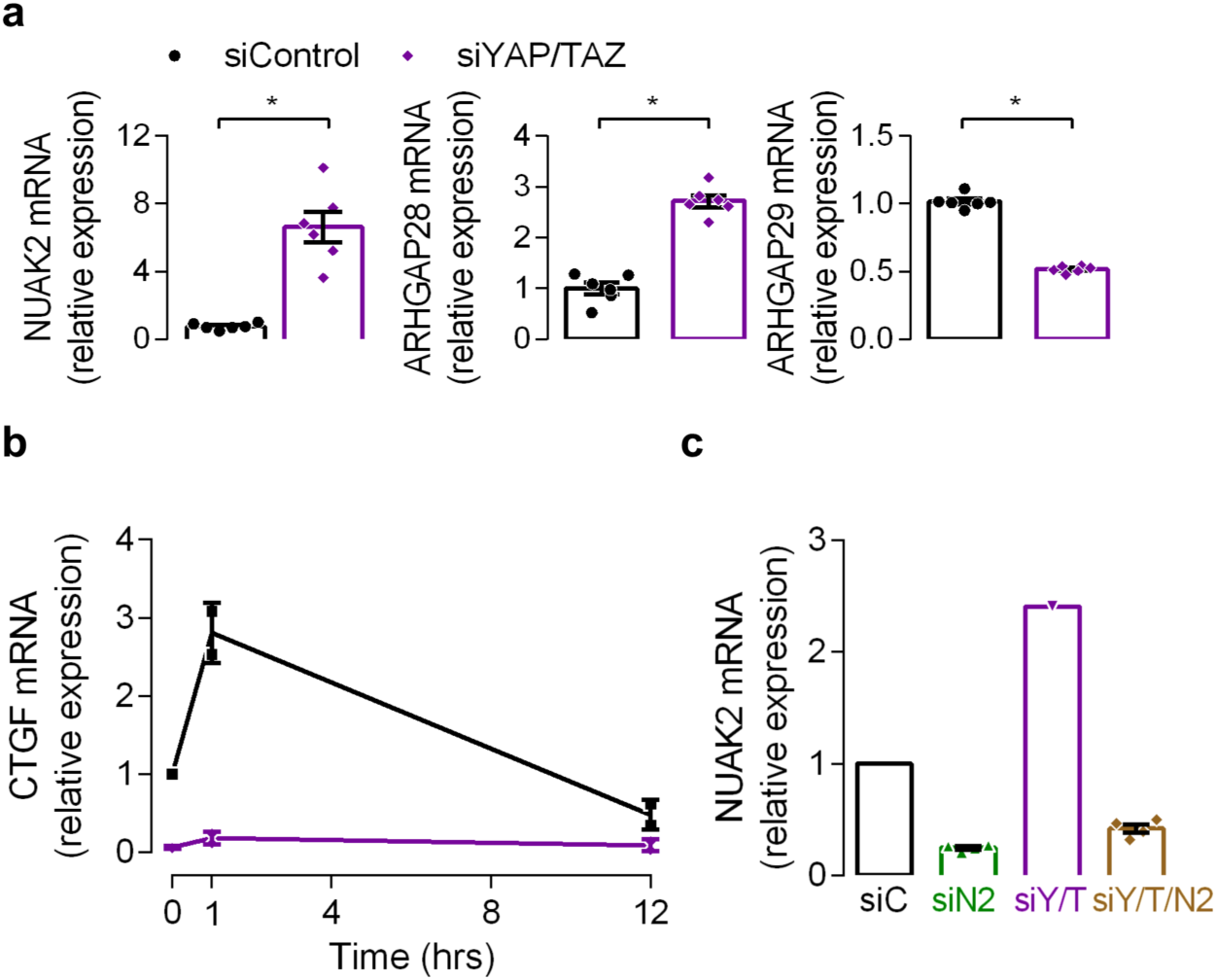
YAP/TAZ depletion decreases CTGF and ARHGAP29 expression while increasing NUAK2 and ARHGAP28 expression in ECFCs. Confluent ECFCs were scratched and allowed to migrate for either 0, 1, or 12 hours then lysed and RNA isolated for RT-qPCR. **a**, ECFCs were lysed 1 hour after initiation of migration and screened for YAP/TAZ regulated cytoskeletal regulators NUAK2, ARHGAP28, and ARHGAP29, n = 6, * p < .0001, two-tailed Student’s unpaired t-test. **b,** CTGF expression during migration, n = 2. **c,** Confluent ECFCs depleted of YAP/TAZ and/or NUAK2 were lysed and RNA used for RT-qPCR, n = 4. Summary statistics are represented as mean ± s.e.m.

**Supplementary Figure 7:**
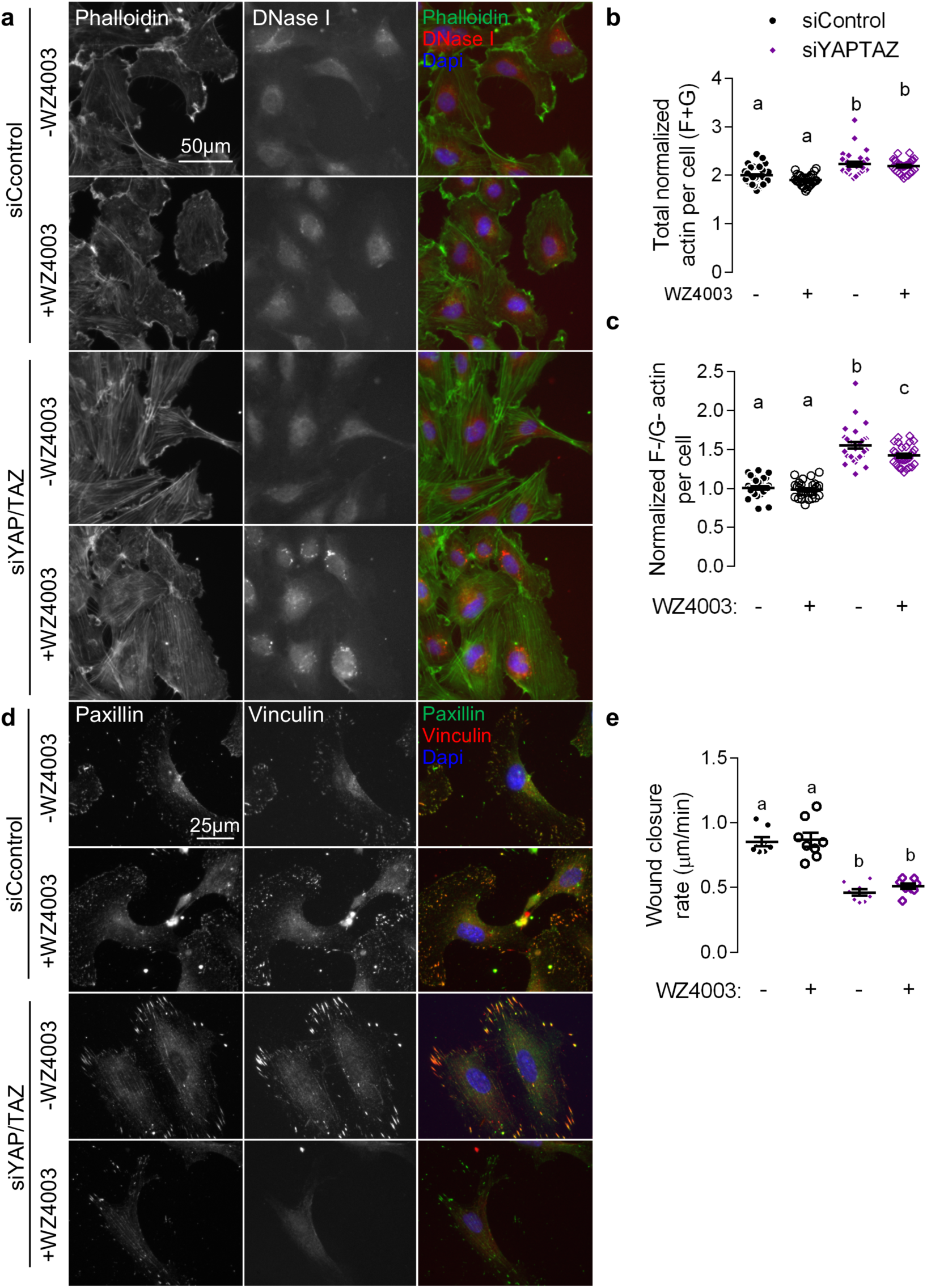
NUAK2 inhibition with WZ4003 partially restores cytoskeletal defects in actin polymerization and focal adhesion morphology, but not migration in YAP/TAZ depleted cells. Migrating ECFCs were depleted of YAP/TAZ and treated with 3µM WZ4003, the NUAK2 inhibitor, for 8 hours then fixed for immunofluorescence. **a,** Representative images of F- and G-actin visualized with Alexa Fluor 594-conjugated phalloidin and Alexa Fluor 488-conjugated DNase I. **b,** Total actin measured as the sum of normalized phalloidin and DNase I intensity. n = 30 cells, p < 0.006, ANOVA with Tukey’s *post hoc* test. **c,** F-actin fraction measured as phalloidin / DNase I intensity, p < 0.04, ANOVA with Tukey’s *post hoc* test. **d**, Cells were treated as described above and then triton-extracted concurrent with fixation for immunofluorescence and analysis of structural focal adhesions. Representative images of vinculin and paxillin visualized with 594- and 488-conjugated secondary. **e,** Confluent ECFCs were scratched and imaged over 12 hours and wound closure rate quantified. n = 5-8, p < 0.0001, ANOVA with Tukey’s *post hoc* test.

**Supplementary Figure 8:**
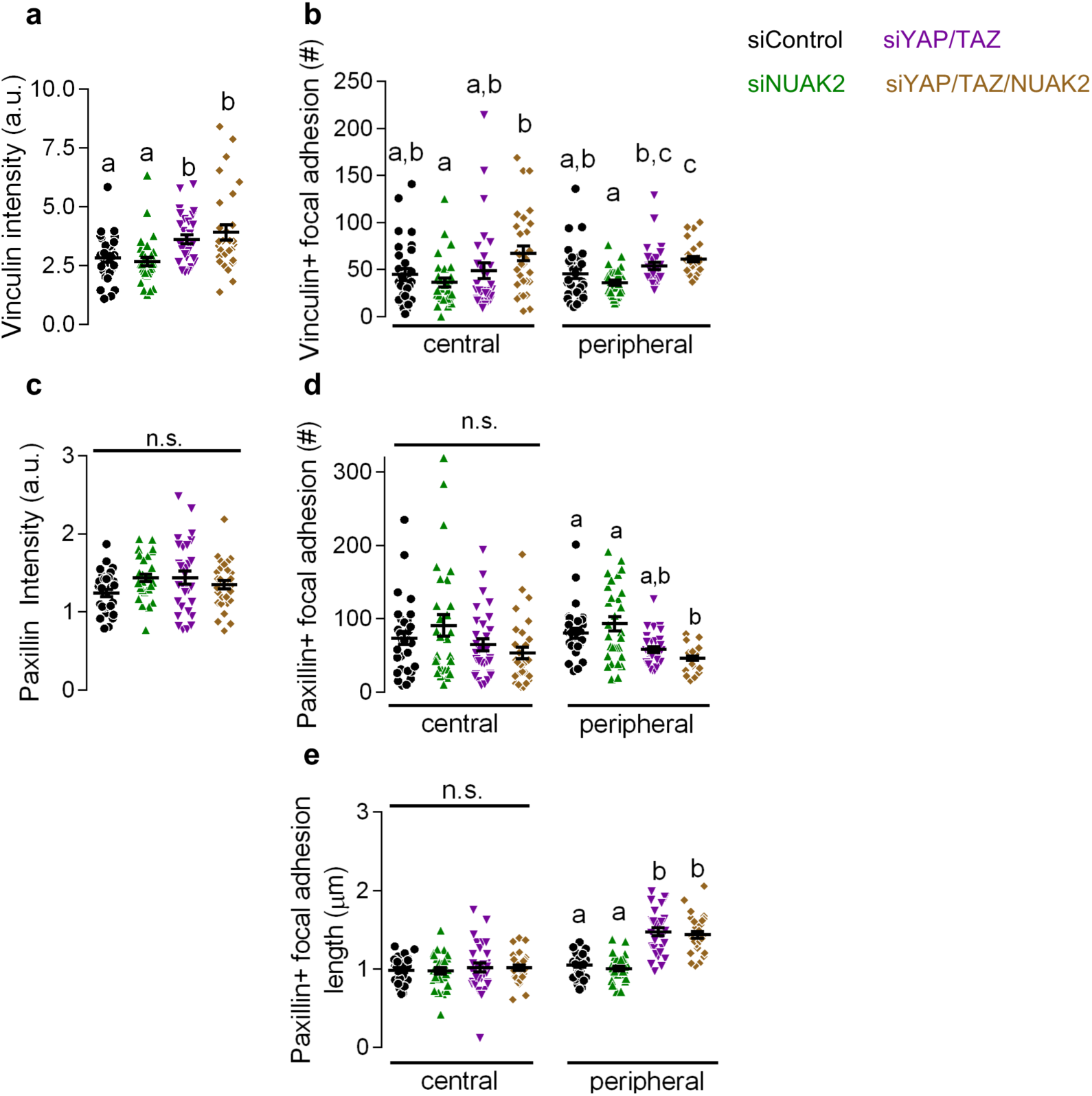
NUAK2 enhances focal adhesion maturation, but does not significantly affect paxillin incorporation into focal adhesions. ECFCs depleted of YAP/TAZ and/or NUAK2 were plated on collagen coated glass coverslips for 24 hours then triton-extracted concurrent with fixation for immunofluorescence and analysis of structural focal adhesions. **a,** Whole cell structural vinculin intensity. n = 30 cell, p < 0.02, ANOVA with Tukey’s *post hoc* test. **b,** Individual cell compartments were subdivided into peripheral and central (5 µm from the edge of the cell) regions to detect spatial differences in focal adhesion morphology (cf. Figure 9b). Focal adhesion length (cf. Figure 9c), an indicator of maturation, and number, an indicator of formation, where found in individual cells at both peripheral and central regions. Central (p > 0.01) and peripheral (p < 0.02) vinculin+ focal adhesion number. **c,** Whole cell structural paxillin intensity. n = 30 cell, p > 0.1, ANOVA with Tukey’s *post hoc* test. **d,** Central (p > 0.08) and peripheral (p < 0.002) paxillin+ focal adhesion number. ANOVA with Tukey’s *post hoc* test. **c,** Central (p > 0.9) and peripheral (p < 0.0001) paxillin+ focal adhesion length. ANOVA with Tukey’s *post hoc* test. Summary statistics are represented as mean ± s.e.m.

**Supplementary Figure 9:**
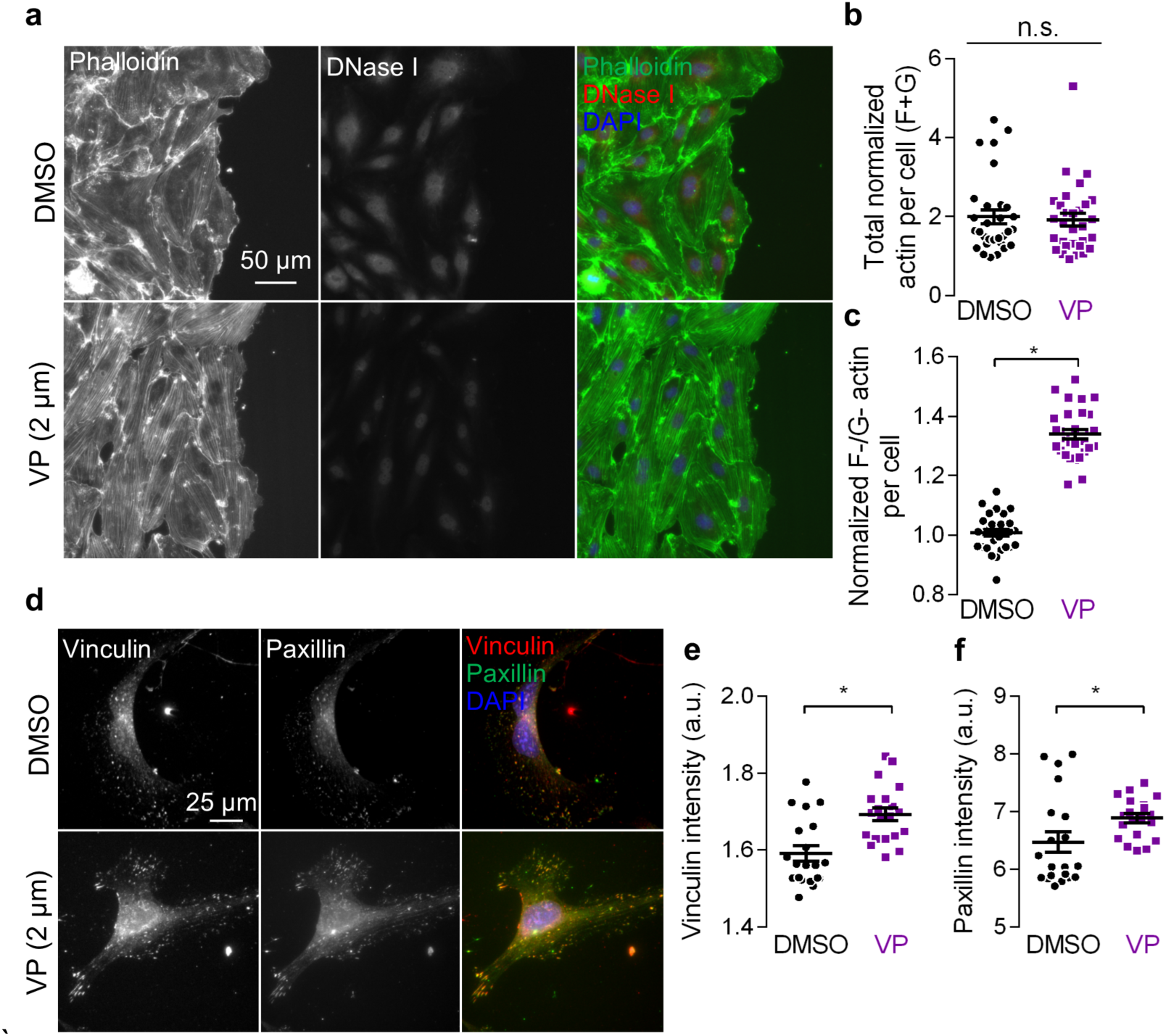
Inhibition of the YAP/TAZ-TEAD interaction with verteporfin phenocopies the cytoskeletal defects observed in YAP and TAZ depleted cells. Migrating ECFCs were treated with 2 µM verteporfin, the YAP/TAZ-TEAD interaction inhibitor, for 8 hours then fixed for immunofluorescence. **a,** Representative images of F- and G-actin visualized with Alexa Fluor 594-conjugated phalloidin and Alexa Fluor 488-conjugated DNase I. **b,** Total actin measured as the sum of normalized phalloidin and DNase I intensity, n = 30 cells, p > 0.78, two-tailed Student’s unpaired t-test. **c,** F-actin fraction measured as normalized phalloidin / DNase I intensity. p < 0.0001, two-tailed Student’s unpaired t-test. **d,** Cells were treated as described above and then triton-extracted concurrent with fixation for immunofluorescence and analysis of structural focal adhesions. Representative images of vinculin and paxillin visualized with 594- and 488-conjugated secondary, respectively. **e,f,** Single cell fluorescence intensity of vinculin (**e)** and paxillin (**f)** incorporation into structural focal adhesions. n = 30, p > 0.0009, two-tailed Student’s unpaired t-test. Summary statistics are represented as mean ± s.e.m.

**Figure s10:**
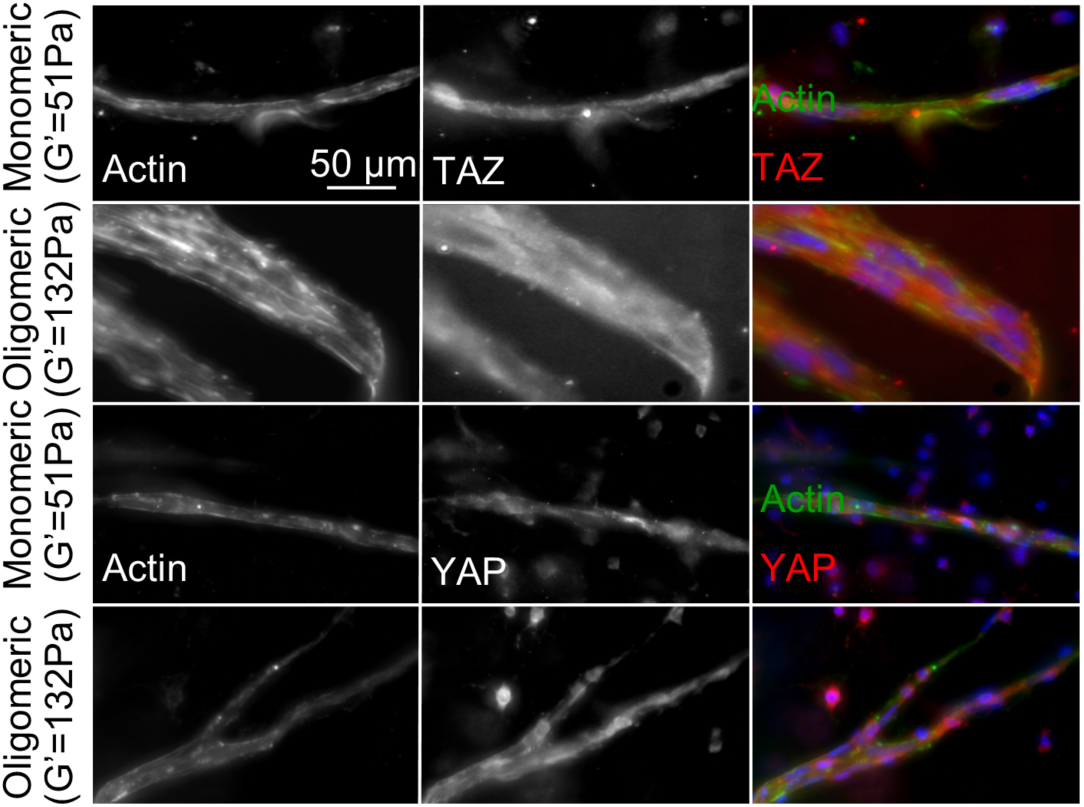
Oligomeric collagen matrices promote angiogenic sprouting from aortic rings. Aortic ring explants were embedded in oligomeric (G’ = 132 Pa) or monomeric (G’ = 51 Pa) collagen matrices. 7 days after explants were embedded whole matrices with explants were fixed for immunofluorescence. Actin and YAP or TAZ were visualized with 488-conjugated phalloidin or 594-conjugated secondary, respectively, nuclei were visualized by DAPI.

